# C24 sphingolipids play a surprising and central role in governing cholesterol and lateral organization of the live cell plasma membrane

**DOI:** 10.1101/212142

**Authors:** K. C. Courtney, W Pezeshkian, R Raghupathy, C Zhang, A Darbyson, J. H. Ipsen, D. A. Ford, H. Khandelia, J. F. Presley, X Zha

**Affiliations:** Department of Biochemistry, Microbiology & Immunology, University of Ottawa, 501 Smyth Road, Ottawa, Ontario, K1H 8L6, Canada.; MEMPHYS, Center for Biomembrane Physics, University of Southern Denmark, Campusvej 55, Odense 5230 M, Denmark; Chronic Disease Program, Ottawa Hospital Research Institute, 501 Smyth Road, Ottawa, Ontario, K1H 8L6, Canada.; Edward A. Doisy Department of Biochemistry and Molecular Biology, Saint Louis University School of Medicine, 1100 South Grand Blvd, St. Louis, MO, 63104, USA.; Department of Anatomy and Cell Biology, McGill University, 3640 rue University, Montreal, H3A 0C7, Canada.

## Abstract

Mammalian cell sphingolipids, primarily with C24 and C16 acyl chains, reside in the outer leaflet of the plasma membrane. Curiously, little is known how C24 sphingolipids impact cholesterol and membrane microdomains. Here, we generated giant unilamellar vesicles and live mammalian cells with C24 or C16 sphingomyelin exclusively in the outer leaflet and compared microdomain formation. In giant unilamellar vesicles, we observed that asymmetrically placed C24 sphingomyelin suppresses microdomains. Conversely, C16 sphingomyelin facilitates microdomains. Replacing endogenous sphingolipids with C24 or C16 sphingomyelin in live HeLa cells has a similar impact on microdomains, characterized by FRET between GPI-anchored proteins: C24, but not C16, sphingomyelin suppresses submicron domains in the plasma membrane. Molecular dynamics simulations indicated that, when in the outer leaflet, the acyl chain of C24 sphingomyelin interdigitates into the opposing leaflet, thereby favouring cholesterol in the inner leaflet. We indeed found that cholesterol prefers the inner over the outer leaflet of asymmetric unilamellar vesicles (80/20) when C24 sphingomyelin is in the outer leaflet. However, when C16 sphingomyelin is in the outer leaflet, cholesterol is evenly partitioned between leaflets (50/50). Interestingly, when a mixture of C24/C16 sphingomyelin is in the outer leaflet of unilamellar vesicles, cholesterol still prefers the inner leaflet (80/20). Indeed, in human erythrocyte plasma membrane, where a mixture of C24 and C16 sphingolipids are naturally in the outer leaflet, cholesterol prefers the cytoplasmic leaflet (80/20). Therefore, C24 sphingomyelin uniquely interacts with cholesterol and governs the lateral organization in asymmetric membranes, including the plasma membrane, potentially by generating cholesterol asymmetry.

**Statement of Significance:** The plasma membrane bilayer of mammalian cells has distinct phospholipids between the outer and inner leaflet, with sphingolipids exclusively in the outer leaflet. A large portion of mammalian sphingolipids have very long acyl chains (C24). Little is known how C24 sphingolipids function in the outer leaflet. Mutations in the ceramide synthase 2 gene is found to decrease C24. This severely perturbs homeostasis in mice and humans. Here, we investigated unilamellar vesicles and mammalian cells with C24 sphingomyelin exclusively in the outer leaflet. We provide evidence that outer leaflet C24 sphingomyelin suppresses microdomains in model membranes and live cells by partitioning cholesterol into the inner leaflet. We propose that C24 sphingolipids are critical to the function of the plasma membrane.

## Introduction

Lateral membrane microdomains, or lipid rafts, have long been regarded as a fundamental feature of the plasma membrane in mammalian cells (1). These domains are thought to be phase-separated from the more fluid environment of the bilayer membrane, primarily through spontaneous side-by-side associations of sphingolipids and cholesterol. In support of this hypothesis, micron-sized domains have been observed in giant unilamellar vesicles (GUVs) and giant plasma membrane vesicles (GPMVs) (2–4). Nevertheless, with the exception of transient nanodomains (5, 6), micron-sized domains have not been visualized in live mammalian cells. This has been attributed to protein/lipid interactions with the cytoskeleton and/or the complex membrane heterogeneity (7). Nevertheless, in contrast to most model membranes, a unique feature of the plasma membrane of live mammalian cells is their phospholipid asymmetry. One notable example is the asymmetrical distribution of sphingolipids, which reside almost exclusively in the outer leaflet (8, 9). Physiological sphingolipids primarily have two acyl chains, C16 and C24 (10). Our current understanding of cholesterol-sphingolipid interactions are primarily derived from model membranes made with C16 or C18 sphingomyelin (SM) in both leaflets, in contrast to physiological membranes (3, 11, 12). With their very long acyl chains, C24 sphingolipids have been observed to interact with cholesterol distinctly from C16 sphingolipids (13), perhaps even more so in asymmetric membranes. Indeed, replacing C24 with C16 sphingolipids due to a mutation in ceramide synthase 2 (CerS2) was recently shown to result in metabolic defects in mouse models. In humans, genome-wide associations between similar mutations and metabolic syndrome have also been reported (14, 15). The precise mechanism for these defects is not known presently. However, it does suggest the critical importance of C24 sphingolipids physiologically. In contrast to C16 sphingolipids, C24 sphingolipids have a unique tendency for transbilayer interdigitation in bilayer membranes, due to the significant mismatch in length between the acyl chain and sphingosine backbone (16). Here, we investigated how C24 SM interacts with cholesterol in asymmetric membranes and its impact on the lateral organization of GUVs and also on the plasma membrane in live mammalian cells.

## Results

### C24 SM suppresses microdomains in GUVs

We first examined the effect of C24 or C16 SM (milk or egg SM) on microdomain formation in three-component GUVs, a widely used model of the plasma membrane (3). Until recently, GUVs were only made of symmetric membranes (3). Without cholesterol, symmetric GUVs composed of DOPC^*^/DPPC or DOPC/SM did not produce stable micron-sized domains (Fig. 1A, *a-c* & Supplementary Fig. 1). Introducing cholesterol into these symmetric GUVs led to phase separation, which can be visualized by the DPPE probes, with NBD-DPPE (green) in the liquid ordered phase (L_o_) and rhodamine-DPPE (red) in the lipid disordered phase (L_d_) (Fig. 1A, *d-f* & Supplementary Fig. 1). Cholesterol-containing symmetric GUVs with either C24 SM or C16 SM formed similar microdomains (Fig. 1A, *e* & *f*) that were initially small but gradually coalesce into large domains (Supplementary ***movie 1***). Thus C24 and C16 SM behaved similarly in symmetric membranes in terms of microdomain formation.

**Figure 1.**
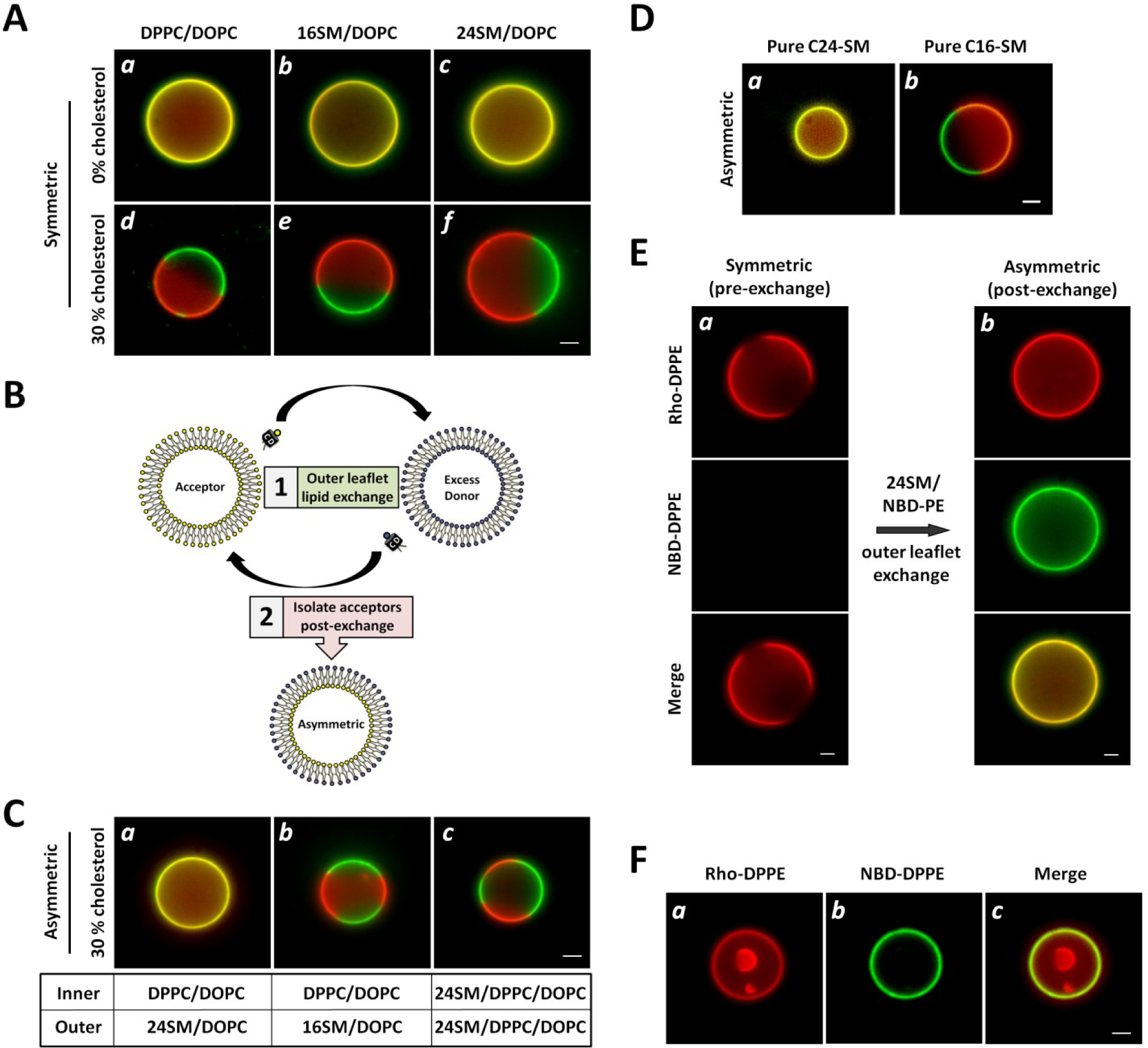
Very long acyl chain SM abolishes optically resolvable microdomains when placed exclusively in the outer leaflet of GUVs. **A**) In symmetric GUVs, cholesterol was necessary for the phase separation of saturated and unsaturated phospholipids, including GUVs with very long acyl chain SM (C24 SM) in both leaflets. Symmetric vesicles were visualized by incorporation of 0.05% rhodamine-DPPE and NBD-DPPE during electroformation. **B**) Pictogram of outer leaflet phospholipid exchange. Symmetric GUVs are converted to asymmetric GUVs by incubation with HP-αCD and excess donor lipid. After exchange of outer leaflet lipids, the acceptor GUVs are isolated from the donors and HP-αCD by filtration, resulting in asymmetric GUVs. **C**) (***a***) In asymmetric GUVs, introducing C24 SM in the outer leaflet of DPPC/DOPC/cholesterol vesicles suppressed visible microdomain formation. (***b***) Asymmetric vesicles with C16 SM in the outer leaflet formed microdomains. (***c***) Symmetric C24 SM/DPPC/DOPC/cholesterol (scrambled) GUVs formed microdomains. **D**) Pure synthetic C16 and C24 SM produced the same result as natural C16 (egg) and C24 (milk) SM. **E**) (***a***) Prior to incorporation of C24 SM, DPPC/DOPC/cholesterol GUVs display microdomains and lack NBD-DPPE. (***b***) Microdomains in DPPC/DOPC/cholesterol GUVs disappeared after incorporation of C24 SM and NBD-DPPE into the outer leaflet during outer leaflet exchange. **F**) Encapsulated symmetric DPPC/DOPC/cholesterol GUVs remained phase-separated inside asymmetric GUVs with exofacial C24 SM. Images were captured at 20 °C and were representative of a homogeneous population of 50-100 vesicles. Experiments were independently verified at least 3 times. Asymmetric vesicles were visualized by initially labelling acceptor vesicles with 0.05% rhodamine-DPPE, followed by incorporation of NBD-DPPE during outer leaflet lipid exchange. Scale bar: 5µm.

We next produced asymmetric GUVs by an exchange protocol (Fig. 1B) (17–19), taking advantage of the fact that phospholipids flip-flop slowly (t_1/2_ is usually days). This method can place either C24 or C16 SM exclusively in the outer leaflet (9). Remarkably, introducing C24 SM into the outer leaflet of GUVs completely abolished the microdomains (Fig. 1C, *a*) at a wide range of temperatures (3–37 °C) and cholesterol concentrations (0-50%) (Supplementary Fig. 2A & 2B). C16 SM, when similarly placed into the outer leaflet, persistently promoted microdomains (Fig. 1C, *b*), as in the symmetric membranes above (Fig. 1A, *e*). Moreover, if C24 SM asymmetry was destroyed by generating GUVs with equivalent lipid compositions as in Fig. 1C, *a*, except C24 SM were now in both leaflets, microdomains were formed again (Fig. 1C, *c*). Furthermore, pure synthetic C24 SM in the outer leaflet abolished microdomains in asymmetric GUVs, but not synthetic C16 SM (Fig. 1D), clarifying that impurities in the natural C16 (egg) and C24 (milk) sphingomyelin played no significant role. Thus, C24 and C16 SM functions distinctively when placed asymmetrically in GUVs.

To further confirm that C24 SM in the outer leaflet is necessary to abolish the domains, we documented the following events. First, before outer leaflet lipid exchange, all symmetric acceptor GUVs (DPPC/DOPC/cholesterol) with rhodamine-DPPE (red) presented visible microdomains and donor-associated NBD-DPPE (green) was absent (Fig. 1E, *a*). Upon exchange (most rhodamine-DPPE in the outer leaflet is lost), acceptor GUVs acquired NBD-DPPE along with C24 SM from donors to the outer leaflet and, simultaneously, visible microdomains disappeared (Fig. 1E, *b*). Secondly, GUVs occasionally had smaller unilamellar vesicles inside (Fig. 1F, *a*). These encapsulated unilamellar vesicles did not have direct access to donor lipids and thus could not acquire NBD-DPPE (no green) or C24 SM (Fig. 1F, *b*). They retained their visible microdomains after exchange, even though their envelope GUV no longer had microdomains (Fig. 1F, *a*). Together, our data demonstrated a surprising role of C24 SM: it abolishes micron-sized domains, but only when placed exclusively in the outer leaflet of GUVs. SM with shorter acyl chains, such as C16 SM, continued to promote domains even when similarly present only in the outer leaflet of GUVs.

### C24 SM also suppresses sub-micron domains in the plasma membrane of HeLa cells

We next asked whether C24 sphingolipids can similarly influence lateral organization in the plasma membrane of mammalian cells. For this, we performed FRET between CFP-and YFP-GPI-anchored proteins (APs) in live HeLa cells. These CFP and YFP GPI-APs do not form specific molecular-molecular interactions, i.e. dimerization or oligomerization (5). However, they both prefer L_o_ domains in model membranes and the live cell plasma membrane (20). Therefore, if L_o_ domains formed in the plasma membrane, both CFP and YFP GPI-APs should become enriched in these micro or submicron domains. Under this scenario, FRET efficiency would be increased, relative to randomly distributed proteins. We first performed a theoretical simulation of FRET between randomly distributed CFP and YFP, which indicated to us that FRET could be detected in the range of protein density achievable in cultured cells and that FRET efficiency increases linearly with the density of acceptors in this density range (Fig. 2A). To experimentally test this, we co-expressed CFP-and YFP-GPI-APs in HeLa cells, followed by cholesterol depletion to generate a condition where the GPI-APs were known to be randomly distributed (21). We compared the experimental FRET data with the theoretical simulation. We found that the experimental FRET fit well within the simulation data and was mostly linear within this range of protein densities (Fig. 2A, inset, blue dots).

**Figure 2.**
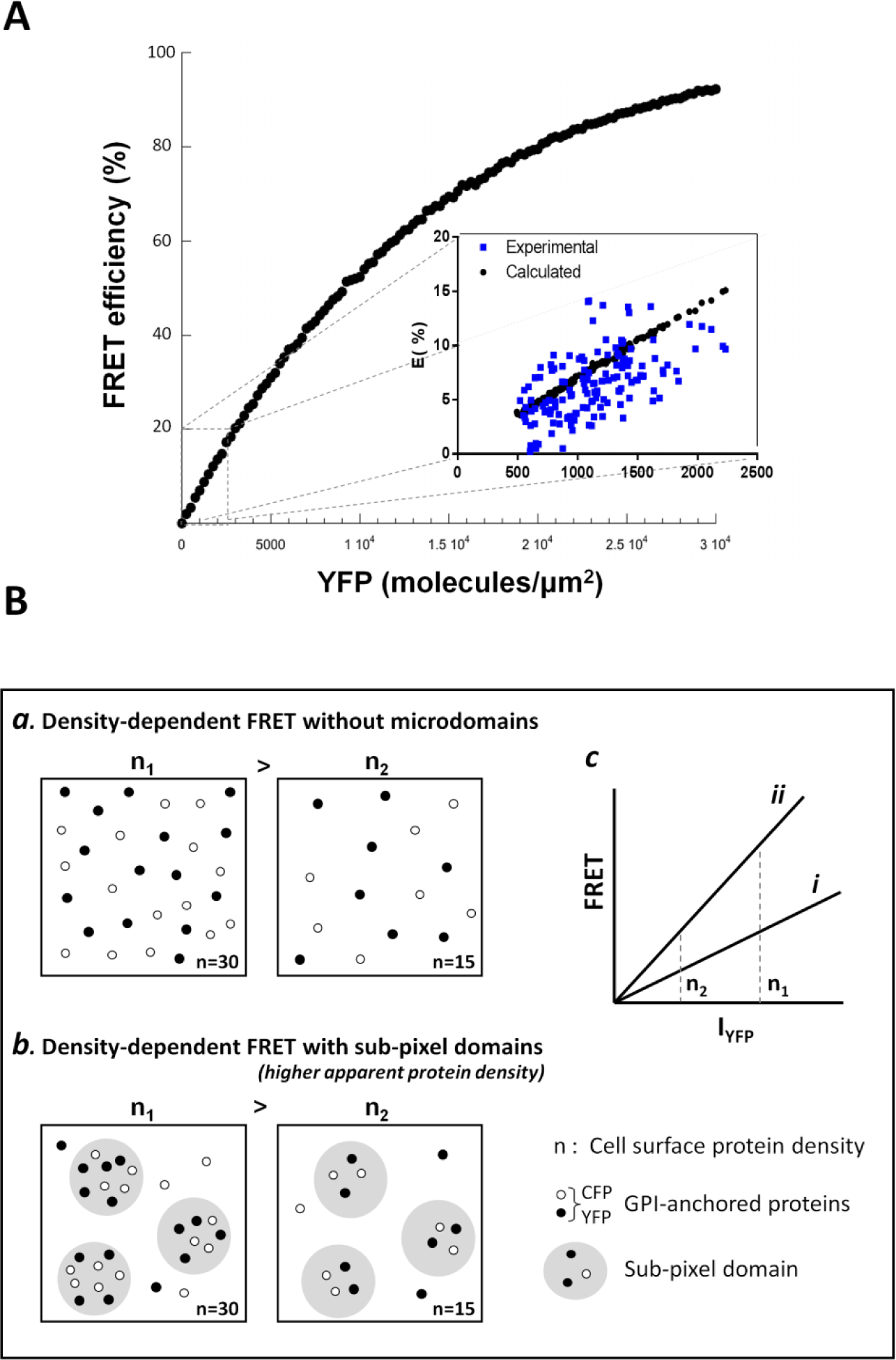
FRET efficiency increases with increasing molecular density of fluorescent proteins, which can monitor relative recruitment of GPI-APs into membrane domains. **A**) Density-dependent FRET efficiency simulation: the solid points represent simulated density-dependent FRET predicted from density of YFP protein. Blue dots represent experimentally obtained FRET efficiencies from the cells treated with saponin. Notice that the stimulated line is linear within the range of the experimental YFP density. **B**) Pictogram of how the presence of membrane domains would affect density-dependent FRET between mCFP and mYFP GPI-APs. GPI-APs prefer L_o_ (submicron domains), which increases the fluorescent protein density within the submicron domains and enhances the FRET.

Our current understanding is that plasma membrane domains are too small to be optically resolved (22). However, the recruitment of CFP and YFP GPI-APs into submicron domains would increase the local density of the GPI-APs and, therefore, enhance the FRET efficiency between CFP- and YFP-GPI-APs. Thus, we envisioned that: (**a**) in the absence of domains: proteins are randomly distributed in the plasma membrane (Fig. 2B, *a*). The dependence of FRET on acceptor concentration (YFP) would be similar to the simulation (Fig. 2B, ***c**, line **i***), which is insensitive to any treatment that abolishes microdomains (i.e. cholesterol depletion); (**b**) in the case where submicron domains are present; CFP- and YFP-GPI APs become concentrated within L_o_ domains (Fig. 2B, *b*). This enhanced recruitment into submicron domains causes CFP- and YFP-GPI APs to be in closer proximity and, therefore, increases FRET efficiency (Fig. 2B, ***c**, line **ii***), compared to randomly distributed proteins (*line **i***). Also, the enhanced FRET should be highly sensitive to cholesterol depletion, as it will disperse submicron domains and put the GPI-anchored proteins into a random distribution, as in *line **i***.

To understand how sphingolipid acyl chain length influences microdomains, we generated HeLa cells with C16 SM or C24 SM in the outer leaflet of the plasma membrane. This was achieved by first depleting all sphingolipids with myriocin and fumonisin b1 (M+F), which inhibits SM biosynthesis and ceramide synthase activity (23, 24). The cells were then replenished with C16 or C24 SM using SM/γ-CD complexes. As shown in Fig. 3A, native HeLa cells (DMSO) have both C16 and C24 sphingolipids with C24 sphingolipids being the most abundant. M+F depleted nearly all sphingolipids; subsequent supplementation successfully replenished the plasma membrane with C16 SM or C24 SM (Supplementary Fig. 3). Cells were viable and the replenished sphingolipids were indeed correctly inserted into the outer leaflet of the plasma membrane, verified by sphingomyelinase (SMase) induced endocytosis. SMase is known to hydrolyze sphingolipids in the outer leaflet of the plasma membrane and hence create an imbalance in the surface area between two leaflets, leading to spontaneous endocytosis (25), shown in control cells here (Supplementary Fig. 4). As expected, in cells depleted sphingolipids (M+F), SMase was not able to induce endocytosis. Critically, cells treated with M+F but subsequently supplemented with C16 or C24 SM were able to be trigged by SMase to endocytose (Supplementary Fig. 4). This demonstrates the availability of SM on the outer leaflet of the plasma membrane. Thus, we concluded that we generated live cells with C16 or C24 SM in the outer leaflet of the plasma membrane, analogous to the asymmetric GUVs in Fig. 1B.

**Figure 3.**
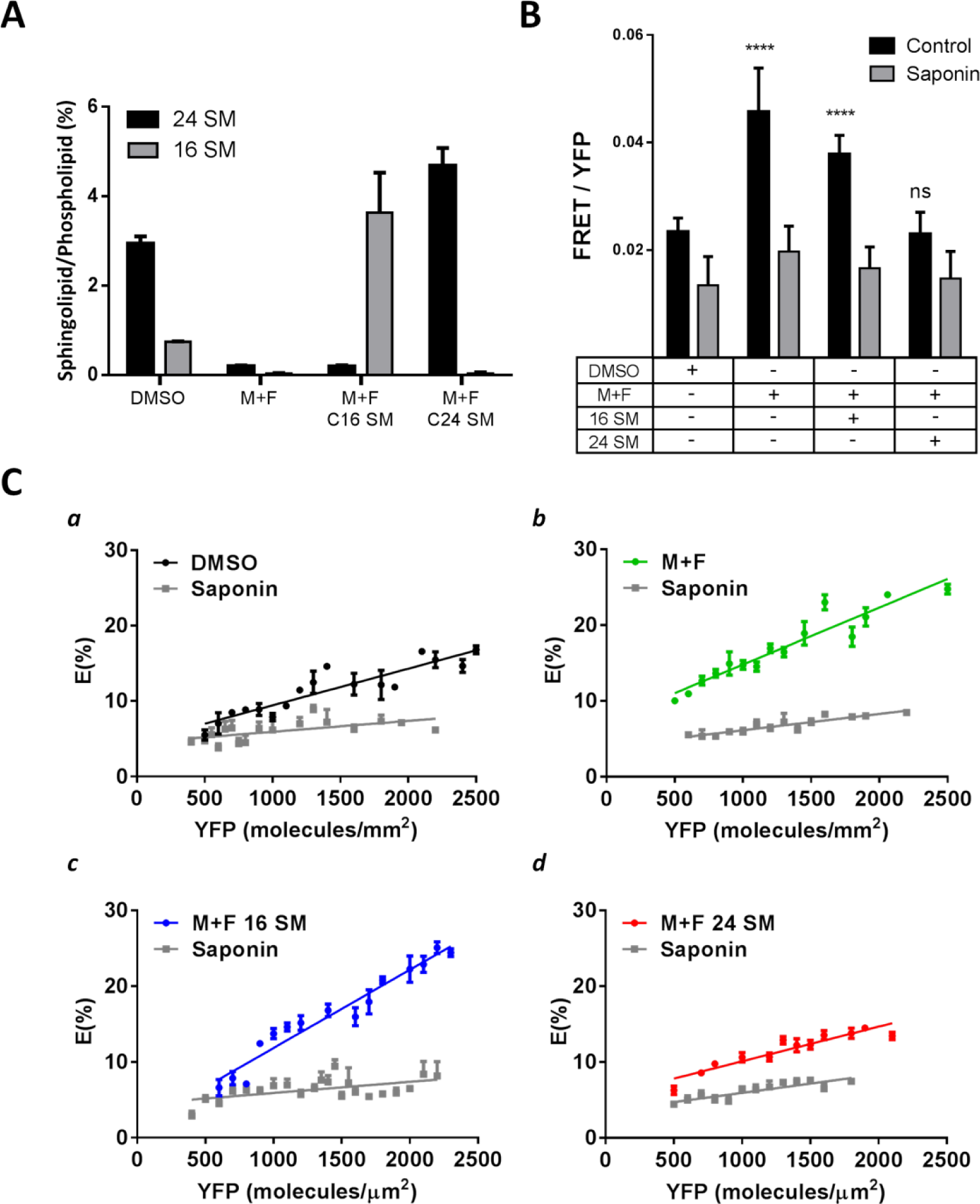
Outer leaflet C24 sphingomyelin suppresses membrane submicron domains in the live cell plasma membrane. **A)** Quantification of sphingolipid acyl chain lengths determined by thin layer chromatography (TLC). Quantities of sphingolipids were determined relative to total cell phospholipid levels by densitometry. Untreated cells (DMSO) were compared to sphingolipid deletion by myriocin and fumonisin b1 (M+F) and subsequent supplementation with C16 or C24 SM. **B)** The effect of cholesterol depletion by saponin on FRET between mCFP and mYFP GPI-anchored proteins in live HeLa cells from sphingolipid depletion and subsequent supplementation with C16 or C24 SM. **C**) FRET efficiency, E (%), between mCFP and mYFP GPI-anchored proteins in live HeLa cells with sphingolipid depletion and subsequent supplementation with C16 or C24 SM.

CFP- and YFP-GPI-APs were then expressed in the HeLa cells. The CFP- and YFP-GPI-APs were primarily localized on the plasma membrane (Supplementary Fig. 5), as reported previously (6, 20). We observed the highest FRET efficiency in cells replenished with C16 SM (Fig. 3B). Also, sphingolipid depleted cells (M+F) had similarly high FRET efficiency. This is consistent with our earlier observation in asymmetric GUVs: microdomains can form without SM or with C16 SM (Fig. 1A, *d* & *e*). In contrast, FRET efficiency is much less in the control cells (DMSO), which naturally have abundant C24 sphingolipids (Fig. 3A). Moreover, in contrast to cells with C16 SM, the cells that were replenished with C24 SM had low FRET efficiency, identical to the control cells. Importantly, cholesterol depletion decreased FRET most dramatically in C16 SM replenished cells and in sphingolipid depleted cells (M+F) (Fig. 3C, *b* & *c*), again in line with the existence of cholesterol-rich L_o_ submicron domains in these cells prior to cholesterol depletion. At the same time, cholesterol depletion produced little change in FRET in either C24 SM replenished or control cells (Fig. 3C, *a* & *d*), indicative of limited submicron domains in the native plasma membrane or the plasma membrane with C24 SM. Therefore, in the plasma membrane of untreated HeLa cells, submicron domains are less prominent than in cells without native sphingolipids or with C16 SM in the outer leaflet. Conversely, replenishment of C24 SM into the outer leaflet diminished the plasma membrane submicron domains and returned the cells to their native state.

We found that we had to express the GPI-AP at a relatively high density (~1000/µm^2^) to achieve reliable density-dependent FRET. However, this density is less than 0.1% of membrane molecules (26), which is well within the range for labelling model membranes with fluorescent phospholipid analogues (2). Furthermore, only C24 SM, not C16 SM, returned FRET to the level of untreated cells (Fig. 3C, *a* & *d*). It is thus plausible that the FRET experiments reported here reflect changes in submicron domains in the plasma membrane.

### Molecular dynamics (MD) simulations and potential of mean force (PMF) suggest interdigitation of C24 SM and potential impact on cholesterol

The interaction between cholesterol and SM is believed to be essential for membrane microdomain formation (7, 27). However, asymmetrically placed C24, but not C16, SM unexpectedly abolished microdomains in GUVs (Fig. 1) and suppressed submicron domains the live cell plasma membrane (Fig. 3). This led us to speculate that C24 SM may interact with cholesterol differently from C16 SM. Indeed, all-atom molecular dynamics (MD) simulations and potential of mean force (PMF) calculations on asymmetric bilayers with either C16 or C24 SM in the outer leaflet (Supplementary Fig. 6) demonstrated several striking differences (Fig. 3A & 3B and Supplementary Fig. 7–9). First, cholesterol favours the inner leaflet (5.5 k_B_T less free energy) if C24 SM is in the outer leaflet (Fig. 4A, a and Supplementary Fig. 7), whereas the SM containing leaflet is preferred by cholesterol if C16 SM is in the outer leaflet (Fig. 4B, *a*). Secondly, the energy barrier for cholesterol to flip from the outer to the inner leaflet is significantly smaller when C24 SM is in the outer leaflet (6 k_B_T), compared to that of C16 SM (15 k_B_T). The effect of C24 SM was equivalent when cholesterol was moving from the inner to outer leaflet or outer to inner leaflet (Supplementary Fig. 8). Thus, energetically, cholesterol would favour the inner leaflet when C24 SM in the outer leaflet. Moreover, the atom density profile indicates that the C24 SM acyl chain from the outer leaflet significantly penetrates into the inner leaflet of the bilayer (Fig. 4A, *b*, *red peak*); while 16 SM is fully contained within the outer leaflet (Fig. 4B, *b*) (Supplementary Fig. 7 for other phospholipids). We initially hypothesized that inner leaflet cholesterol would directly interact with the interdigitated outer leaflet C24 SM acyl chain; however, we found no evidence of trans-bilayer stacking of cholesterol and C24 SM in opposing leaflets from an analysis of bilayer registry and two-dimensional density maps of cholesterol and SM in the bilayer plane (data not shown). Alternatively, we postulated that C24 SM acyl chain interdigitation would increase the density in the inner leaflet near the centre of the bilayer and induce mechanical instability by perturbing phospholipid packing. To compensate, cholesterol could either move into the inner leaflet to fill the gap and/or C24 SM would be pushed up towards the aqueous phase, which weakens outer leaflet C24 SM-cholesterol interactions. Consistent with this notion, the MD simulations indicated that C24 SM engages in weaker H-bonding with cholesterol in the outer leaflet than C16 SM (Supplementary Fig. 9). Weaker H-bonding was strongly correlated with reduced electrostatic interaction energy between C24 SM and cholesterol (-134 ± 8 kJ/mol), compared to C16 SM (-190 ± 15 kJ/mol). Outer leaflet C24 SM also exhibited higher electrostatic and Lennard-Jones interaction energies with the solvent molecules, compared to C16 SM, confirming that the C24 SM was indeed pushed up towards the aqueous phase in the asymmetric membranes (Supplementary Fig. 9H). Therefore, when C24 SM is in the outer leaflet, reduced H-bond capacity in the outer leaflet could further favour cholesterol in the inner leaflet, although other factors may also be contributing.

**Figure 4.**
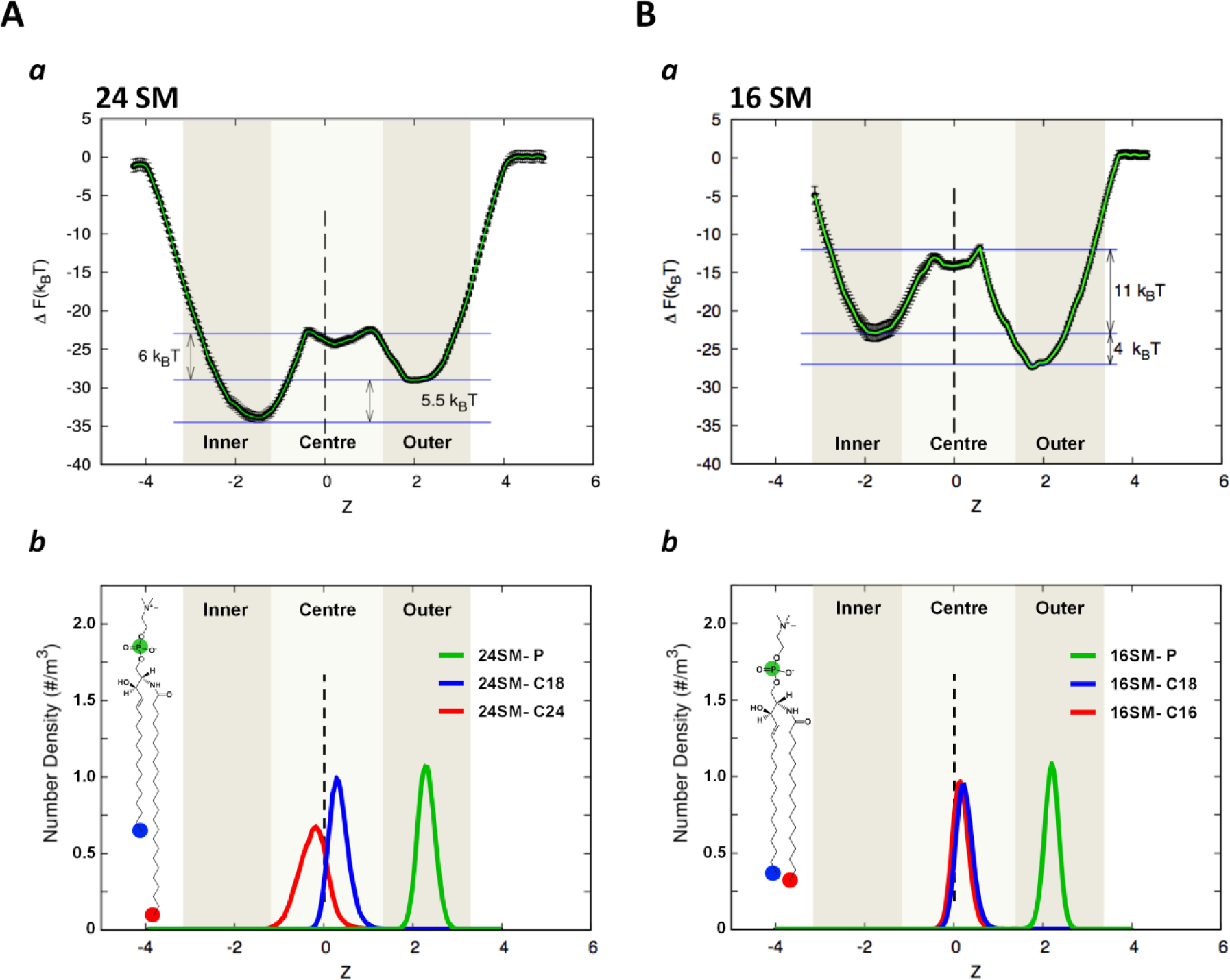
Cholesterol displays a preference for the inner bilayer leaflet when very long acyl chain sphingomyelin is in the outer leaflet. **A) (*a*)** Free energy profile of transferring a cholesterol molecule from outer leaflet to the inner leaflet in a C24 SM asymmetric membrane shows cholesterol prefers the inner leaflet. The Z-axis refers to the z-distance between the position of the pulled cholesterol molecule relative to the center of the bilayer, Z=0 (see Methods). Errors in the free energy profile were determined using the bootstap analysis method. **(*b*)** Normalized density profile for specific atoms of C24 SM in the C24 SM membrane system. Only the head group phosphate (green) and terminal acyl chain carbons (blue and red) are displayed to demonstrate the depth of acyl chain penetration into the bilayer. **B) (*a*)** Free energy profile for transferring a cholesterol molecule from outer leaflet to the inner leaflet in a C16 SM asymmetric membrane shows cholesterol has a slight preference for the outer leaflet. The Z-axis and errors are the same as in A. **(*b*)** Normalized density profile for specific atoms of C16 SM in the C16 SM membrane system. Only the head group phosphate (green) and terminal acyl chain carbons (blue and red) are displayed to demonstrate the depth of acyl chain penetration into the bilayer.

### Development of a novel protocol to quantify cholesterol in each leaflet of large unilamellar vesicles

We next proceeded to experimentally test the effect of C24 SM on cholesterol partitioning. Unlike phospholipids, cholesterol flip-flops rapidly between leaflets (t_1/2_ < sec) (11), which has greatly hindered the analysis of cholesterol partitioning in bilayer membranes. In order to quantify cholesterol partitioning, it was necessary to develop an experimental protocol to prevent cholesterol flip-flopping. Cholesterol was observed by electron spin resonance (ESR) to stop rotating within phospholipid membranes at 0 ºC (28). It was also reported that cholesterol movement, either within phospholipid membranes or between membranes, were prevented at 0 ºC regardless of membrane phospholipid compositions (29). Interestingly, methyl-β-cyclodextrin (MCD), a high affinity cholesterol chelator, was seen to remove cholesterol from unilamellar vesicles at low temperature, albeit at slower rate (30). We postulated that MCD could specifically remove cholesterol from the outer leaflet of unilamellar vesicles at 0 ºC, when cholesterol flip-flop was stopped. This concept was tested using symmetric large unilamellar vesicles (LUVs). Phospholipids and cholesterol in symmetric LUVs with diameter 100 nm should be evenly partitioned in each leaflet. Therefore, a valid analysis should detect cholesterol partitioned evenly, i.e. 50/50, between leaflets. Importantly, such 50/50 partitioning should be completely independent of phospholipid compositions of the LUVs.

We then carried out all the experiments at 0 ºC under extremely stringent temperature control: all experiments were performed in the cold room in an ice-water bath (0 ºC). In addition, all the utensils were pre-cooled in ice water so that no temperature change occurred during sample handling (see method for further detail). We first confirmed the unilamellar nature of LUVs (100 nm) (supplementary Fig. 10A). The LUVs used here only contain a trace amount of cholesterol (0.01%), so that the LUVs maintained their structural integrity after cholesterol removal (Supplementary Fig. 10B). It was also noted that, due to a tighter binding of cholesterol at lower temperature, cyclodextrins can sequester cholesterol highly efficiently at low temperature (30).

We then added membrane-impermeable MCD to the medium to extract cholesterol at 0 ºC and 37 ºC, respectively. If no cholesterol is flip-flopping between leaflets, for example at 0 ºC, MCD should extract precisely 50% of the cholesterol from symmetric LUVs, regardless of phospholipid composition. However, if cholesterol is allowed to freely flip-flop, as occurs at 37 ºC, all cholesterol (100%) is accessible to MCD (Fig. 5A). Indeed, this is exactly the case: MCD removed 53% (± 2.9) at 0 ºC and 100% cholesterol at 37 ºC from symmetric LUVs (Fig. 5B), independent of phospholipid compositions, including LUVs with C24 SM (Fig. 5C-E). Furthermore, MCD also extracted precisely 50% cholesterol from symmetric LUVs at -5 ºC (in an ethylene glycol bath) (Fig. 5F). This further confirms that MCD is able to extract cholesterol at 0 and -5 ºC, which is solely dependent upon the sidedness of the cholesterol, rather than interactions with phospholipids. We, therefore, concluded that we established a valid protocol to prevent cholesterol flip-flop, which can be used to quantify cholesterol partitioning in LUVs.

**Figure 5.**
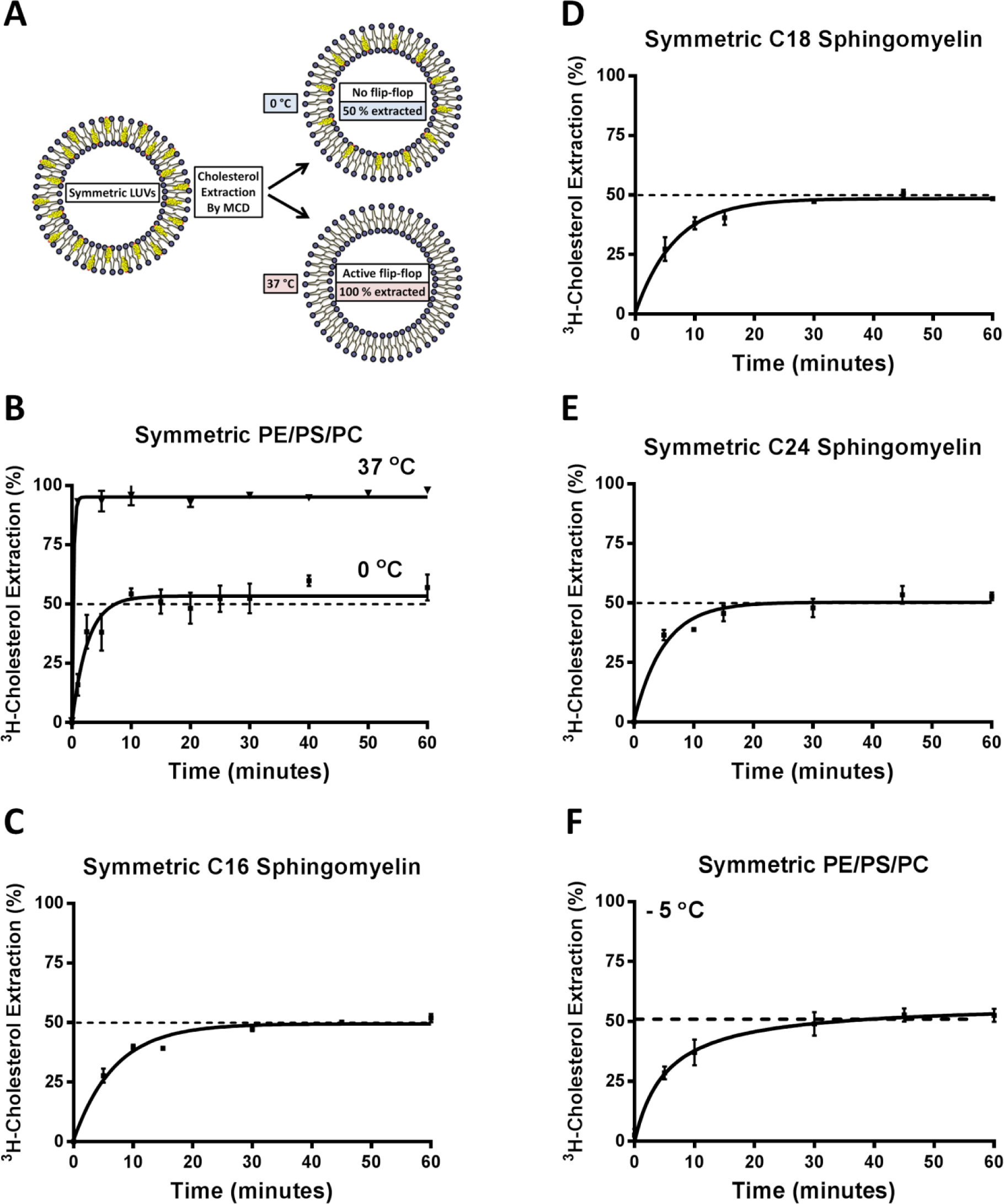
Cholesterol flip-flop is prevented at 0 °C and evenly partitioned between the inner and outer leaflet of symmetric LUVs. **A)** Pictogram of cholesterol extraction from a symmetric membrane bilayer by MCD at 0 °C and 37 °C. Membranes are labelled with trace of 3H-cholesterol and allowed to equilibrate. Lowering temperature to 0 °C prevents cholesterol flip-flop, which facilitates selective outer leaflet cholesterol extraction and quantification by MCD. At 37 °C, cholesterol flip-flop is active and MCD can extract 100 % of the cholesterol due to inner leaflet cholesterol flipping outward. **B**) In symmetric POPC/POPS/POPE (1:1:1) LUVs, MCD was able to extract 100% cholesterol at 37°C but at 0 °C, only 50% cholesterol is removable, verifying that cholesterol flip-flop is prevented. **C**) Cholesterol extraction from symmetric C16 SM LUVs at 0 °C showed cholesterol is evenly partitioned between inner and outer leaflets. **D**) Cholesterol extraction from symmetric C18 SM LUVs 0 °C showed cholesterol is evenly partitioned between inner and outer leaflets. **E**) Cholesterol extraction from symmetric C24 SM LUVs 0 °C showed cholesterol is evenly partitioned between inner and outer leaflets. **F**) Cholesterol extraction from symmetric POPC/POPS/POPE (1:1:1) LUVs -5 °C showed cholesterol is evenly partitioned between inner and outer leaflets.

### C24 SM in the outer leaflet concentrates cholesterol into the inner leaflet in large unilamellar vesicles

We next generated asymmetric LUVs with C24, C16, or C18 SM in the outer leaflet, which were validated by anisotropy and mass spectrometry (Supplementary Fig. 11). Remarkably, when C24 SM was in the outer leaflet, MCD could only maximally remove 20% cholesterol at 0 ºC (Figure 6A). This suggests that 80% of the cholesterol is inaccessible to MCD, i.e. in the inner leaflet. Moreover, if these asymmetric LUVs were dissolved, lyophilized and reformed into symmetric vesicles (C24 SM in both leaflets or “scrambled”), MCD again removed 50% of the cholesterol (Fig. 6A, inset). Thus, the most plausible interpretation is that C24 SM in the outer leaflet caused cholesterol to become partitioned 80/20 between the inner and outer leaflet at 0 ºC.

**Figure 6.**
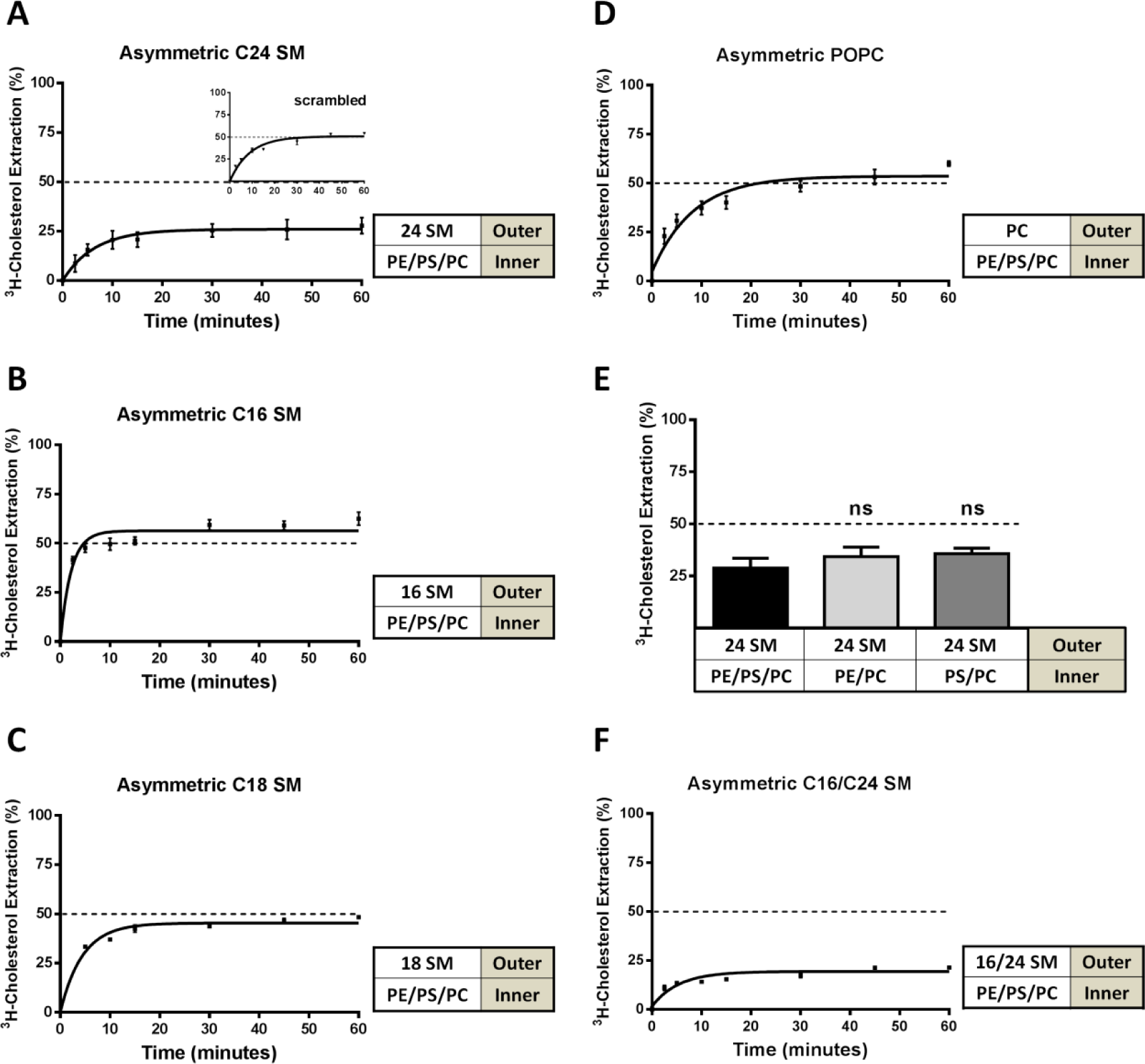
Outer leaflet very long acyl chain sphingomyelin concentrates cholesterol into the inner leaflet. Asymmetric LUVs were generated by outer leaflet lipid exchange and cholesterol was extracted by 5 mM MCD at 0 °C. **A**) In asymmetric LUVs with PC/PE/PS in the inner leaflet, outer leaflet C24 SM shifted cholesterol to the inner leaflet. Abolishing asymmetry by scrambling the asymmetric C24 SM LUVs into symmetric LUVs with an identical composition recovered 50/50 cholesterol partitioning (inset). **B**) Cholesterol extraction from asymmetric C16 SM LUVs at 0 °C shows cholesterol is evenly partitioned between bilayer leaflets. **C**) Cholesterol extraction from asymmetric C18 SM LUVs at 0 °C shows cholesterol is evenly partitioned between bilayer leaflets. **D**) Cholesterol extraction from asymmetric POPC LUVs at 0 °C shows cholesterol is evenly partitioned between bilayer leaflets. **E**) Cholesterol extraction at 0 °C from asymmetric C24 SM LUVs with PC/PE/PS, PC/PE or PC/PS in the inner leaflet shows C24 SM continues to alter the cholesterol partitioning. **F**) Cholesterol extraction at 0 °C from asymmetric LUVs with both C16 and C24 SM in the outer leaflet. Outer leaflet C16/C24 SM (50:50) continues to alter the cholesterol partitioning. Error bars represent standard error of the mean from at least 3 independent experiments.

Asymmetric LUVs with C16 SM, C18 SM (brain SM), or PC in the outer leaflet maintained the 50/50 cholesterol partitioning, as in symmetric LUVs (Fig. 6B-D). Furthermore, in LUVs with C24 SM in the outer leaflet, inner leaflet PE or PS seemed to contribute little to the 80/20 cholesterol partitioning in our experimental setting (Fig. 6E). Perhaps most importantly, when a mixture of both C24 SM and C16 SM (50/50) were placed in the outer leaflet, LUVs still exhibited the 80/20 cholesterol partitioning (Fig. 6F), identical to the experiments with only C24 SM in the outer leaflet. This indicates that C24 SM plays a more dominant role than C16 SM in cholesterol partitioning, which could be of physiological significance. We found that, in most mammalian cells, C24 sphingolipids are a major species, with C16 sphingolipid being less abundant (Supplementary Fig. 12).

### C24 SM, when in the outer leaflet, also concentrates cholesterol in the inner leaflet in cholesterol-rich LUVs

Mammalian plasma membrane has C24, along with C16, sphingolipids naturally in the outer leaflet (10) and is also cholesterol rich (~30-40 %). The MCD extraction protocol above, while sufficient to demonstrate that cholesterol flip-flopping is prevented at 0 ºC, could not be used with cholesterol-rich membranes. Removing large quantities of cholesterol from cholesterol-rich LUVs (30%) would surely compromise membrane integrity. We hence employed an exchange protocol between cholesterol-rich donor and acceptor LUVs, which are identical except the donor LUVs contained a trace amount of ^3^H-cholesterol (11). A low concentration of βCD was used as a shuttle to facilitate cholesterol exchange between donor and acceptor LUVs (100 fold in excess) (Fig. 7A). This system allows cholesterol to exchange between donor and acceptor LUVs without net mass flow, rather than direct extraction by MCD. Donor LUVs were also biotinylated and bound to streptavidin-coated beads to facilitate separation from the acceptor LUVs in the supernatant. The amount of ^3^H-cholesterol that is accessible to exchange (i.e. in the outer leaflet of the donor LUVs at 0 ºC) will be transferred to acceptor LUVs and quantified. The protocol was first validated with symmetric LUVs with 30 % cholesterol. We indeed found that 50 % of the ^3^H-cholesterol was exchangeable at 0 ºC and 100 % at 37 ºC (Fig. 7B). However, if C24 SM was introduced into the outer leaflet, these cholesterol-rich LUVs only had 20% of the ^3^H-cholesterol accessible for exchange at 0 ºC (Fig. 7C), again suggesting that 80 % of the cholesterol was shielded from exchange, consistent with enriched partitioning into the inner leaflet. This result is also correlated with the disappearance of microdomains in asymmetric C24 SM GUVs (Fig. 1C, *a*). Taken together, our observations from both cholesterol-poor and -rich LUVs are consistent with the notion that outer leaflet C24 SM is a sufficient factor to partition cholesterol into the inner leaflet of asymmetric LUVs.

**Figure 7.**
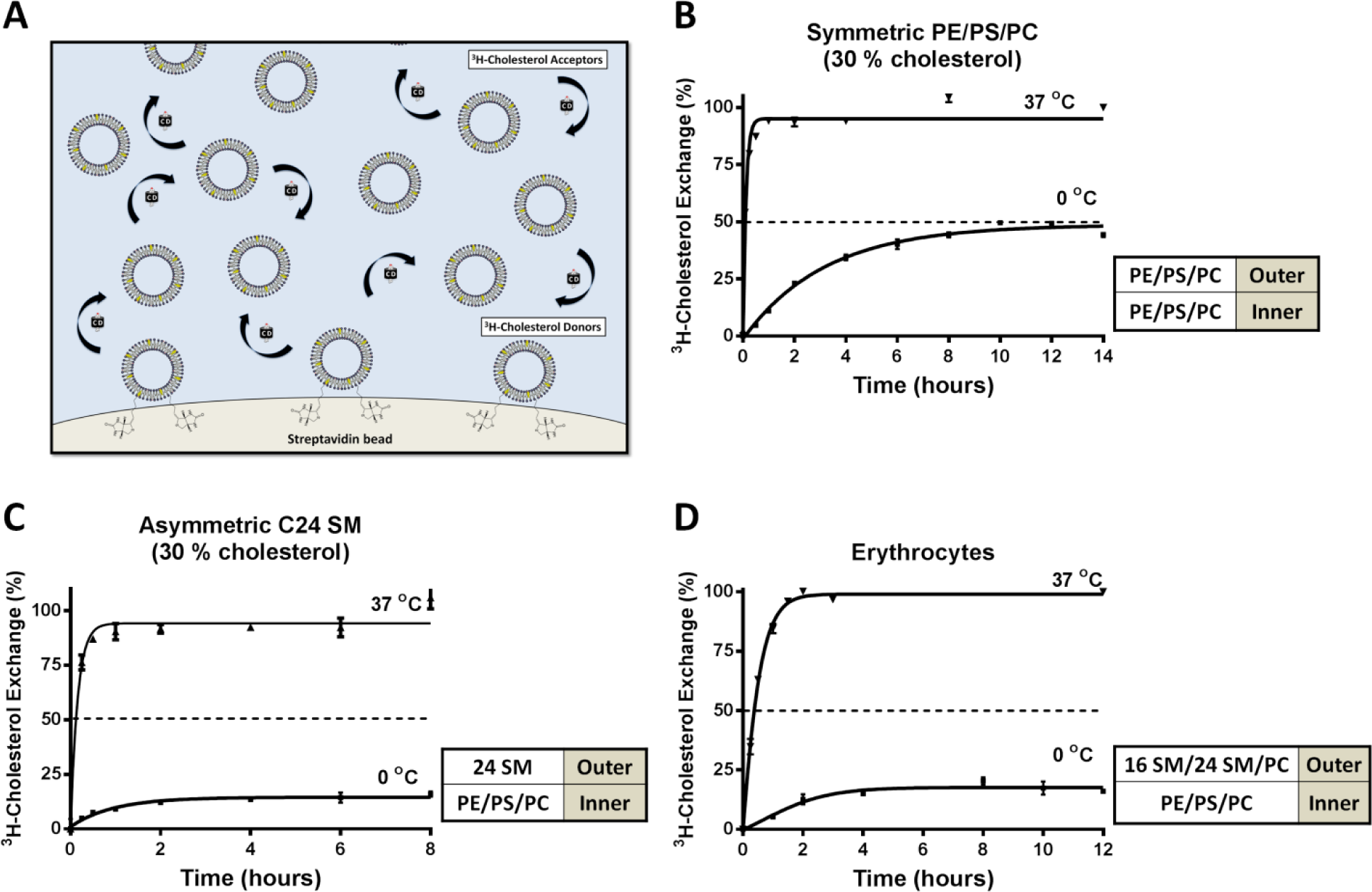
Outer leaflet very long acyl chain sphingomyelin concentrates cholesterol into the inner leaflet of cholesterol-rich LUVs and cholesterol is primarily located in the cytoplasmic leaflet of human erythrocytes. **A**) Pictogram of intermembrane cholesterol exchange facilitated by βCD as shuttle, between donor and acceptor membranes with 30 % cholesterol. ^3^H-cholesterol is transferred from the donor membrane to acceptor membrane (100-fold excess) by βCD and replaced with unlabelled cholesterol. During exchange, the accessible 3H-cholesterol is depleted from the donors, while the total cholesterol content in the donor and acceptor membranes remains unchanged. After exchange, the donor and acceptor populations are separated by brief centrifugation and quantified. **B**) In symmetric POPE/POPS/POPC (1:1:1) LUVs containing 30% cholesterol, cholesterol was evenly distributed between leaflet, as 50% and 100% ^3^H-cholesterol was exchangeable at 0 and 37 °C, respectively. **C**) In 30 % cholesterol LUVs with C24 SM exclusively into the outer leaflet of POPE/POPS/POPC (1:1:1) LUVs, only 20% ^3^H cholesterol was exchangeable at 0 °C and 100% at 37 °C. **D**) In ^3^H-cholesterol labelled human erythrocytes, also only 20% ^3^H cholesterol was exchangeable with at 0 °C with 100 fold unlabelled erythrocytes, demonstrating that cholesterol is primarily in the cytoplasmic leaflet. Error bars represent standard error of the mean from at least 3 independent experiments.

### Live human erythrocytes have substantial amounts of C24 sphingolipids and cholesterol is partitioned 80/20 in the plasma membrane

We next applied the cholesterol exchange protocol to human erythrocytes. Erythrocytes lack internal membranous organelles and are essentially mammalian plasma membranes with predominately C24 sphingolipids (Supplementary Fig. 13). In fact, erythrocytes were instrumental for studies that established phospholipid asymmetry of the plasma membrane, including that of sphingolipids (8, 9, 31). As in LUVs above, we labelled donor erythrocytes with ^3^H-cholesterol. Cells were also biotinylated to be adhered to streptavidin-coated dishes. 100-fold unlabeled erythrocytes were added to the dish as acceptors, again in the presence of βCD. This donor-acceptor system again is preferable over direct cholesterol extraction (32) to minimize perturbing the native cholesterol concentration and avoid haemolysis. We found that only 20 % of the erythrocyte cholesterol was exchangeable at 0 ºC and 100% at 37 ºC (Fig. 7D). The erythrocytes remained intact during the cholesterol exchange (Supplementary Fig. 13C). Thus, in the plasma membrane of erythrocytes, approximately 80% of the cholesterol is shielded from exchange at 0 ºC, consistent with enriched partitioning into the cytoplasmic leaflet. It should be noted that this 80/20 cholesterol partitioning at 0 ºC may not reflect the precise degree of partitioning at physiological temperatures. Nevertheless, only C24 SM in the outer leaflet of asymmetric LUVs produced the 80/20 partitioning at 0 ºC (Fig. 6A and 7C), which is correlated with disappearance of microdomain in GUVs at physiological temperature. The shorter acyl chain SMs did not produce anything other than 50/50 cholesterol partitioning at 0 ºC and did not suppress microdomains in GUVs at physiological temperature. Furthermore, the MD simulations suggest a preferential partitioning of cholesterol into the inner leaflet at 37 ºC, if C24 SM is in the outer leaflet. Intriguingly, studies using fluorescent cholesterol analogs on nucleated mammalian cells similarly found 80/20 partitioning in the plasma membrane at physiological temperature (33, 34).

## Discussion

Taken all observations above together, we propose that C24 sphingolipids, one of the major species of sphingolipids in mammalian cells, have two unique functions on asymmetric membranes, including the plasma membrane. First, when present exclusively in the outer leaflet, C24 sphingolipids most likely enrich cholesterol in the inner or cytoplasmic leaflet (Fig. 6A & 7C). A strong interdigitation into the inner leaflet by the outer leaflet C24 sphingolipids is a potential mechanism (Fig. 4 and Supplementary Fig. 7) (13). Such interdigitation would reduce H-bonding between C24 SM and cholesterol in the outer leaflet and also perturb phospholipid packing in the inner leaflet, generating instability in the membrane (Supplementary Fig. 9). By localizing into the inner leaflet, cholesterol not only fills the potential hydrophobic cavity and thus stabilizes the membrane, but also avoids reduced H-bonding with C24 sphingolipid in the outer leaflet, both of which lead to a more stable state for cholesterol in the inner leaflet (lower energy as shown in Fig. 4A, *a*). Importantly, we observed that, even with equal amounts of C24 SM and C16 SM in the outer leaflet, cholesterol continues to prefer the inner leaflet (Fig. 6F), the state that is energetically more favourable (Fig. 4A, *a* & *b*).

Secondly, asymmetrically placed C24 SM suppresses microdomains in GUVs (Fig. 1). This function is strictly correlated with the preferential partitioning of cholesterol into the inner leaflet of unilamellar vesicles, as shown by cholesterol partitioning experiments (Fig. 6A) and MD simulations (Fig. 4). Our observations by FRET in live cells are also consistent with the notion that C24 SM, naturally in the outer leaflet, limits the formation of submicron domains in the plasma membrane (Fig. 3). We speculate that this could be a consequence of more cholesterol partitioning in to the inner leaflet. In the absence of C24 SM interdigitation (e.g., sphingolipid depletion or C16 SM supplementation), cholesterol asymmetry may be altered in the plasma membrane and promote the formation of submicron domains.

The partitioning of cholesterol in erythrocytes, shown in Fig. 7D, fully supports studies on the partitioning of fluorescent cholesterol analogues, dehydroergosterol and cholestatrienol (33, 34). These studies used a membrane impermeable reagent to collisionally quench the sterol fluorescence from either side of the membrane, thereby fully preserving the native partitioning and flip-flopping dynamics of cholesterol analogues. Unfortunately, most other studies that characterized cholesterol partitioning, or sidedness, relied upon enzymatic reactions or protein probes that form complexes with cholesterol. These approaches would invariably take cholesterol out of their rapidly equilibrium pools and, therefore, perturb the native partitioning of cholesterol (35, 36). In contrast, our 0 ºC protocol was specifically designed not to perturb the native cholesterol distribution. The fact that we reached similar conclusions as the fluorescence quenching studies should also greatly strengthen the conclusion. Moreover, by using native cholesterol, the current study represents another major advance from cholesterol analogues.

Perhaps most importantly, we are able to establish a correlation between the cholesterol partitioning and the formation of microdomains in GUVs. It is attempting to speculate that, the enrichment of cholesterol in the cytoplasmic leaflet of the plasma membrane, as seen in erythrocytes, could create a cholesterol poor outer leaflet. At the same time, the inner leaflet could have a high concentration of cholesterol. Neither of these cases would favour stable microdomains in model membranes (3). This could potentially explain why only transient and nano-scale domains have been reported in live cell plasma membranes. Our FRET study also could only detect minimal submicron domains in control cells.

The cholesterol partitioning or sidedness analysis presented here was carried out using the 0 ºC protocol to essentially “freeze” cholesterol flip-flopping. Our conclusions primarily rely on MCD accessibility to cholesterol in unilamellar vesicles at 0 ºC, which entirely depends on the sidedness of cholesterol. Cholesterol accessibility by MCD is not influenced by phospholipids: all symmetric vesicles with various phospholipid compositions, including those made with C24 SM (Fig. 5E), universally produce precisely 50 % cholesterol extraction by MCD. This is a strong validation of the protocol. It is, therefore, highly unlikely that C24 SM in the outer leaflet of an asymmetric membrane could hinder cholesterol extraction by MCD.

The precise degree of native cholesterol asymmetry in the plasma membrane at physiological temperature remains to be determined. It is possible that the interaction between phospholipids and cholesterol could differ between 0 ºC and physiological temperature. For example, low temperature could potentially cause cholesterol to associate stronger with the inner leaflet lipids, such as PE and PS, compared to outer leaflet SM and PC. As such, cholesterol could become trapped in the inner leaflet when C24 SM is in the outer leaflet. However, only C24 SM was found to disperse microdomains in the asymmetric GUVs and also the submicron domains in live mammalian cells at physiological temperature. Conversely, C16 SM promoted GUV microdomains and submicron plasma membrane domains and was also unable to shift the cholesterol partitioning at 0 ºC. Regardless of the precise degree of cholesterol partitioning, the unique property of C24 SM on microdomains at physiological temperature is correlated with its influence on cholesterol partitioning at 0 ºC. Given the results from the MD simulations and quenching of fluorescent cholesterol analogues, it is likely that outer leaflet C24 SM causes cholesterol to favour the inner leaflet, even at physiological temperature.

Another issue here is the role of the major inner leaflet lipids, PS and PE. PS was found to impact nano-domains in the plasma membrane (21), as well as, the binding of D4 (a perfringolysin O sterol binding domain) to the cytoplasmic leaflet (37). PE was also speculated to influence the partitioning of cholesterol in a bilayer due to effects on membrane curvature (38). However, we did not observe a significant role of PS or PE in LUV cholesterol partitioning (Fig. 6E). Nevertheless, removing these amino-phospholipids in asymmetric LUVs did show a minor trend to reverse the C24 SM effect (Fig. 6E), though failed to reach statistical significance in our experimental system. If PS and PE also favour inner leaflet cholesterol partitioning, this would provide another force to retain cholesterol in the inner leaflet, in addition to the effect of outer leaflet C24 sphingolipids.

Most mammalian cells have a significant amount of C24 along with C16 sphingolipids (Supplementary Fig 12). This suggests that our observations here may have broad physiological implications. Indeed, switching C24 to C16 sphingolipids by changes in CerS2 expression was found to be associated with metabolic syndrome, cancer and neurodegeneration in humans (14, 15, 39, 40). One of the potential consequences of altering the C16/C24 sphingolipid ratio could impact on the native cholesterol sidedness in the plasma membrane, i.e. more cholesterol in the outer leaflet. This would change the dynamics of plasma membrane microdomains and, consequently, protein-protein interactions, contributing to the development of disease.

## Materials and Methods

Phospholipids and liposome accessories were purchased from Avanti Polar Lipids. Cholesterol, trinitrobenzenesulphonic acid (TNBS) and all cyclodextrins used were purchased from Sigma-Aldrich. Radiolabelled (^3^H) cholesterol was purchased from Perkin Elmer. TMA-DPH was purchased from Molecular Probes. Naphtho[2,3-a]pyrene was purchased from Santa Cruz Biotechnologies. Thin layer chromatography plates were acquired from Cedarlane Laboratories. The transfection reagent, Attractene, was purchased from Qiagen. Platinum wire was purchased from Omega Engineering. All other materials and reagents were purchased from Fisher Scientific.

Additional information on materials and methods can be found in Supplementary Information (SI) Materials and Methods.

## Supporting Information

**Supplementary Fig. 1.**
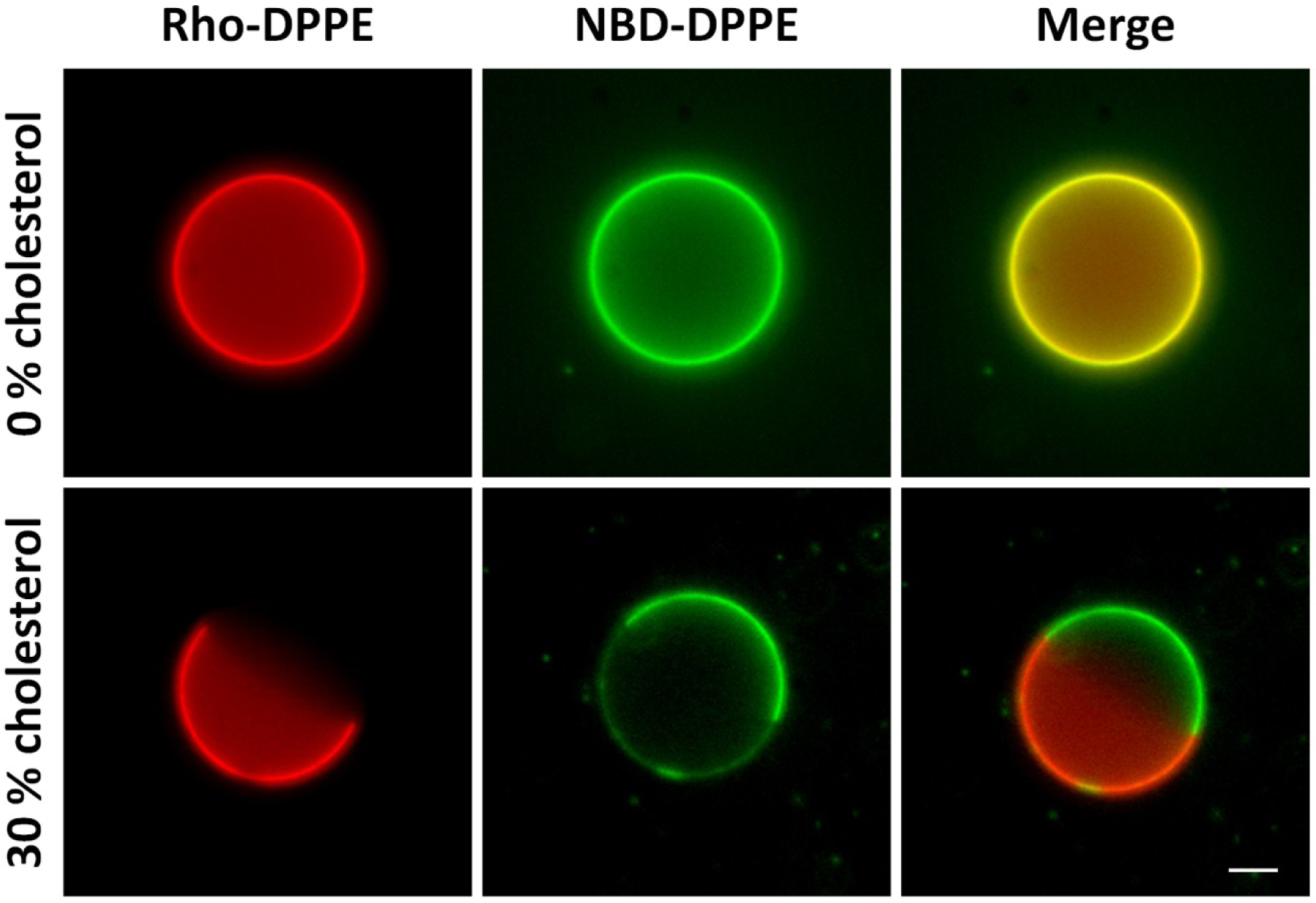
Cholesterol is essential for phase separation in GUVs. Introduction of 30 % cholesterol into GUVs containing saturated DPPC and unsaturated DOPC (1:1) phospholipids causes lipid immiscibility. NBD-DPPE (green) and Rho-DPPE (red) segregate to the liquid ordered (L_o_) and liquid disordered (L_d_) regions, respectively. Scale bar represents 5µm.

**Supplementary Fig. 2.**
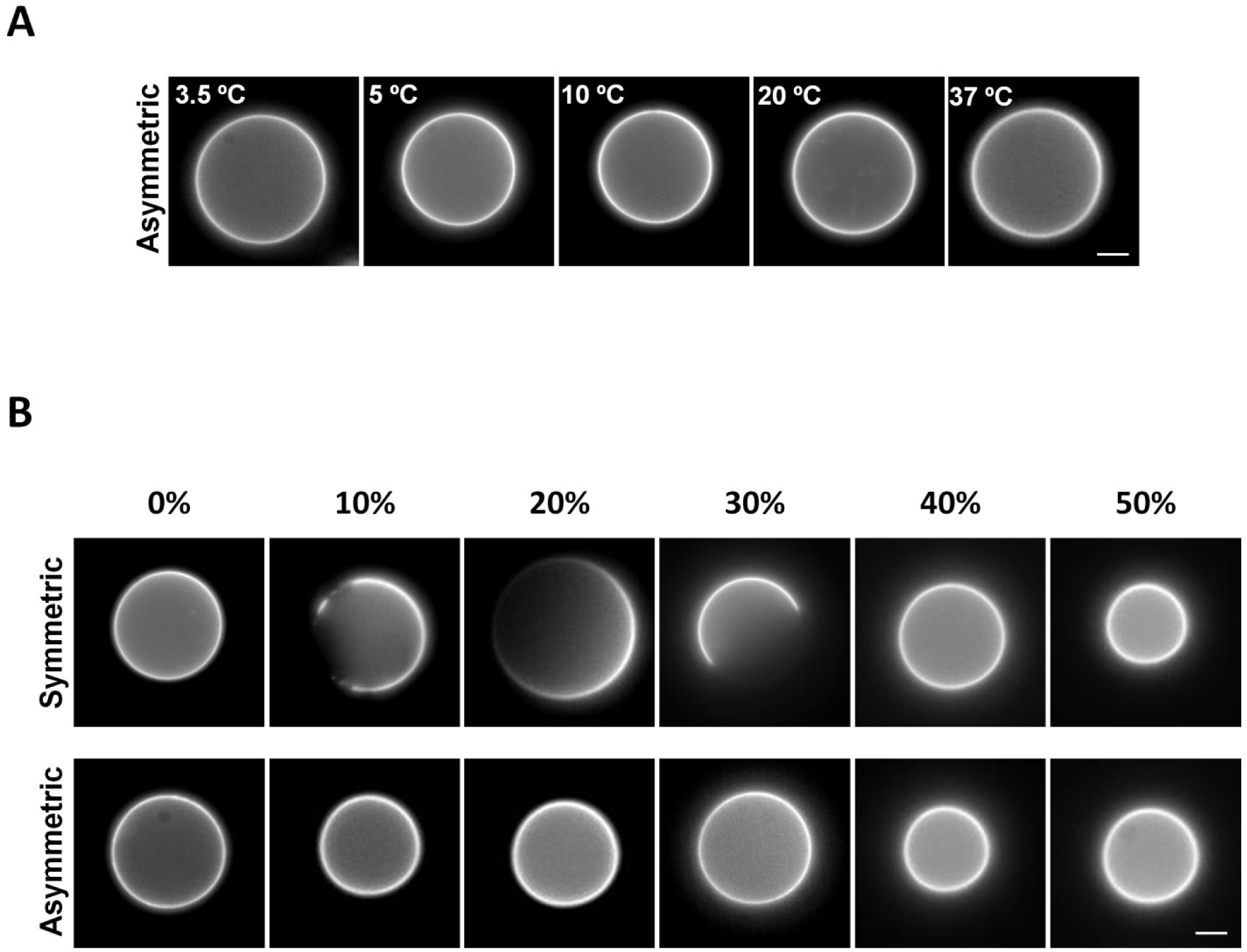
Very long acyl chain sphingolipids in the outer leaflet prevent phase separation in asymmetric vesicles regardless of temperature and cholesterol concentration. **A**) Introduction of C24 SM into the outer leaflet of DOPC/DPPC/cholesterol GUVs prevents microdomain formation in a wide range of temperatures. **B**) Symmetric C24 SM GUVs form membrane domains when the cholesterol concentration is less than 40%. Asymmetric C24 SM GUVs prevent microdomain formation at all cholesterol concentrations. Vesicles were visualized by incorporation of 0.05% rhodamine-DPPE.

**Supplementary Fig. 3.**
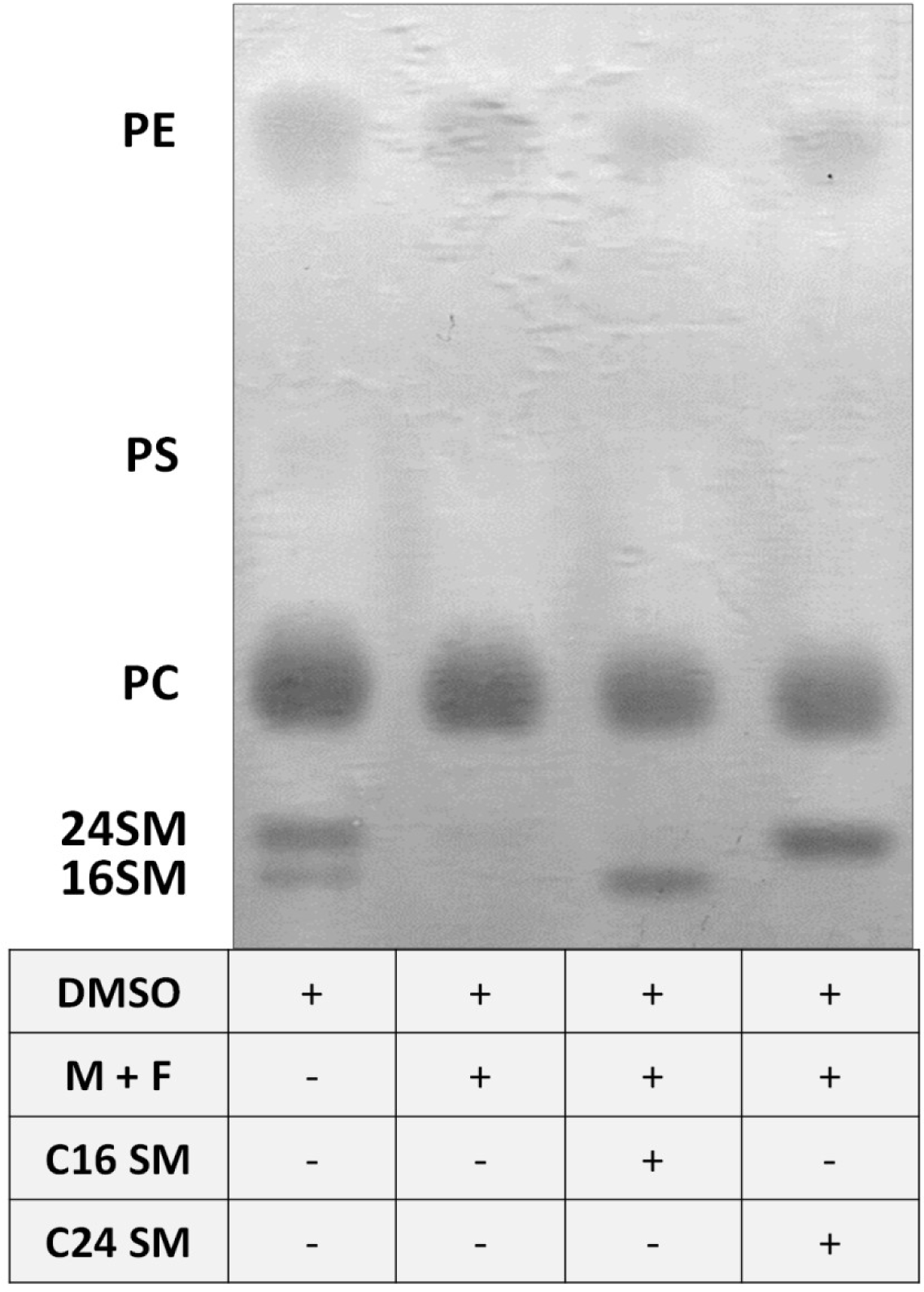
Thin layer chromatography of whole cell lipids after depletion of sphingolipids and subsequent supplementation with C16 or C24 sphingomyelin. HeLa cells were treated with DMSO or myriocin and fumonisin b1 (50 μM) for 3 days. To incorporate the desired sphingolipids, the cells were incubated with a sphingomyelin/γ-cyclodextrin complex containing 10 μM sphingomyelin for 1 hour at 37 °C. The cells were then either characterized by microscopy, or the whole cell lipids were extracted, purified and run on a thin layer chromatography plate. Lipids on the plate were visualized with coomassie blue G stain.

**Supplementary Fig. 4.**
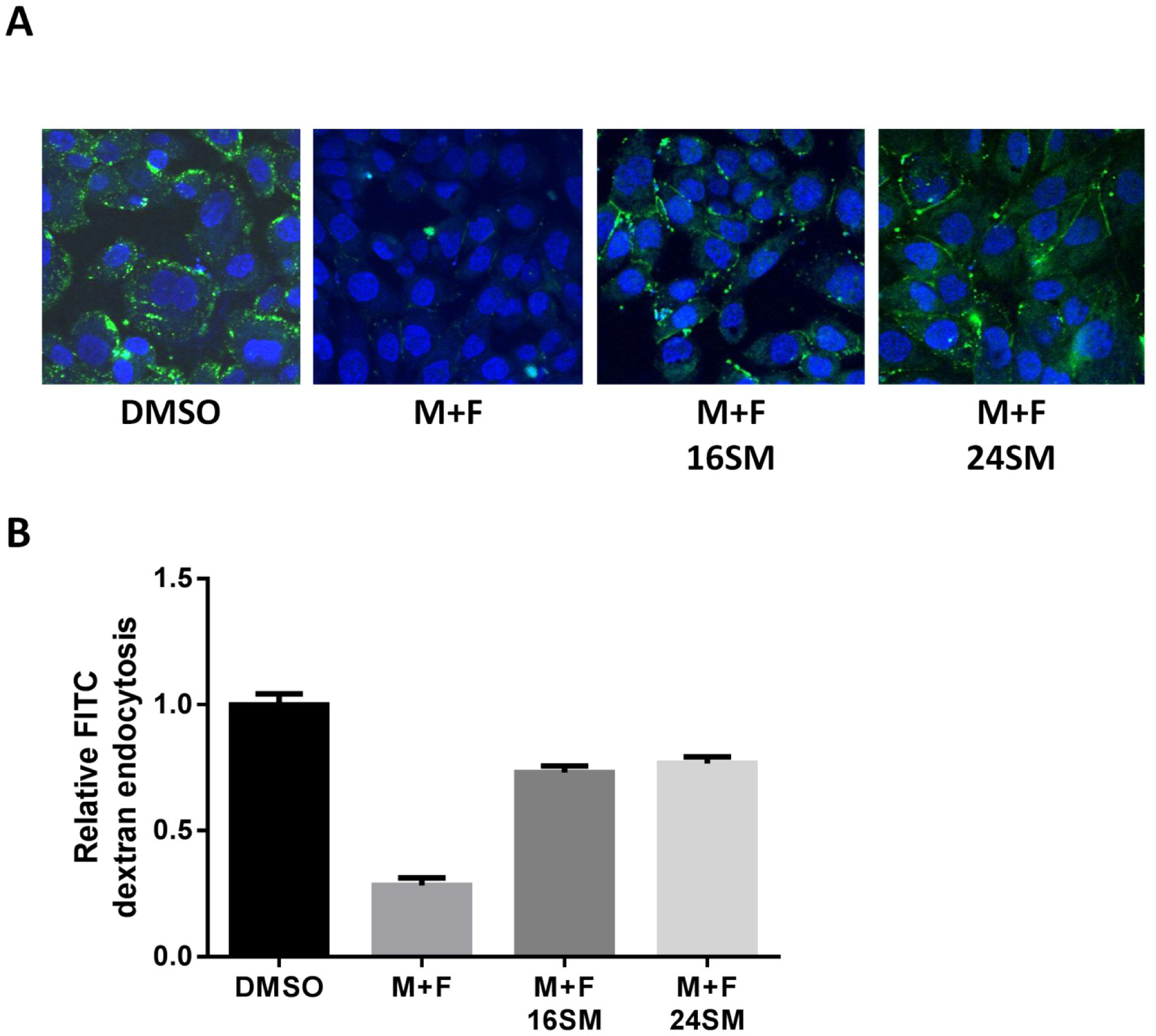
Supplementing HeLa cells with C16 or C24 sphingomyelin restores sensitivity to sphingomyelinase after endogenous sphingolipids have been depleted. **A**) Fluorescence microscopy of FITC-dextran endocytosis in HeLa cells. HeLa cells were treated with either DMSO or Fumonisin B1 and Myriocin for 72 hours. Post drug treatment, C 16 SM or 24 SM complexed to γ-cyclodextrin was added back to the cells for 1 hour. The cells were then washed and incubated in medium containing 5 mg/ml FITC-dextran plus 50 mU/ml sphingomyelinase for 10 min. Cells were fixed and examined by fluorescence microscopy as described in Materials and Methods. **B**) Quantification of endocytosed FITC dextran intensity, relative to the DMSO control. Error bars represent standard error of the mean for three independent experiments.

**Supplementary Fig. 5.**
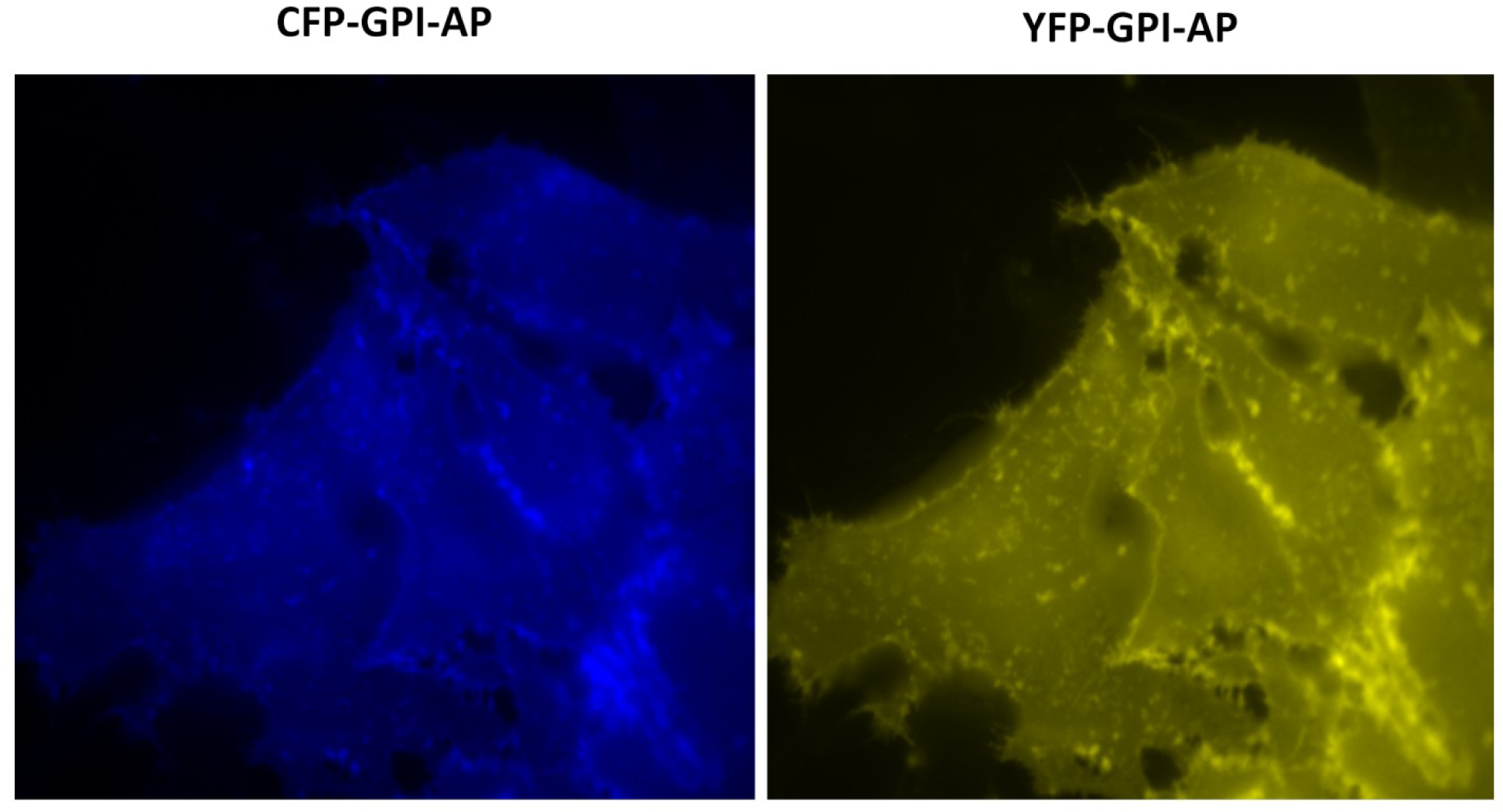
Glycosylphosphatidylinositol anchored proteins (GPI-APs) localize to the plasma membrane in HeLa cells. CFP and YFP GPI-APs were expressed in HeLa cells and visualized by fluorescence microscopy.

**Supplementary Fig. 6.**
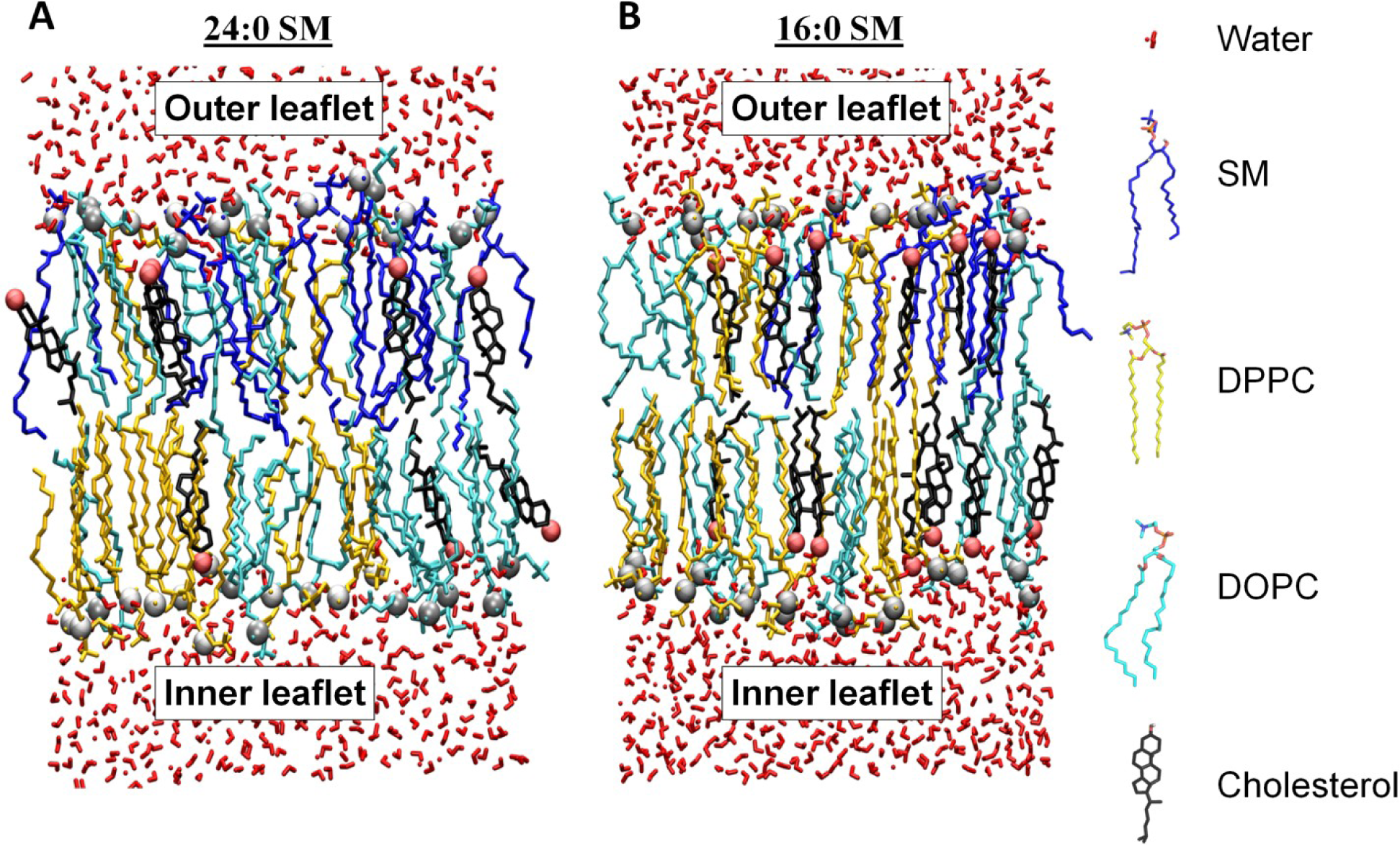
Snapshot of molecular dynamic simulations of asymmetric bilayers with 16:0 and 24:0 SM in the outer leaflet. **A**) Cross sectional image of asymmetric 24:0 SM membrane at the end of the simulation. **B**) Cross sectional image of asymmetric 16:0 SM membrane at the end of the simulation. Lipid composition for both 16:0 and 24:0 SM simulations: 20 SM (blue), 20 DPPC (yellow), 20 DOPC (cyan) and 24 CHL (black) molecules in the upper monolayer and 28 DPPC, 29 DOPC and 24 CHL in the lower monolayer. 5158 water molecules are present. Red and gray spheres are O and P atom of cholesterol and DOPC lipids respectively.

**Supplementary Fig. 7.**
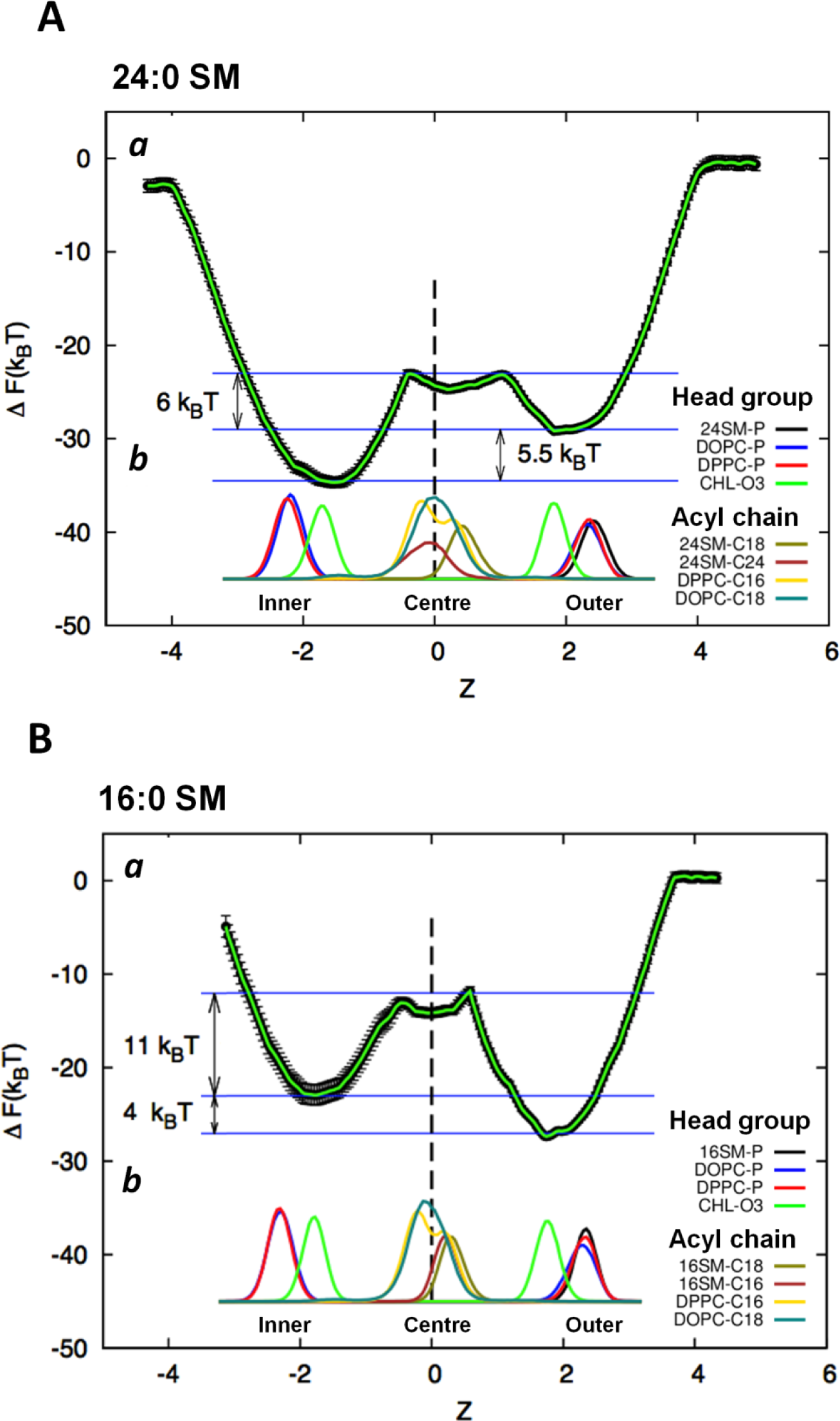
Cholesterol displays an energetic preference for the inner bilayer leaflet when very long acyl chain sphingomyelin is in the outer leaflet. The atomic density of all the lipid species is represented. **A**) (***a***) Free energy profile of transferring a cholesterol molecule from outer leaflet to the inner leaflet in a C24 SM asymmetric membrane shows cholesterol prefers the inner leaflet. The Z-axis refers to the z-distance between the positions of cholesterol relative to the bilayer centre (see Methods). (***b***) Normalized density profile for specific atoms of different lipids in the C24 SM membrane system. The Z-axis refers to the proximity of the lipids relative to the bilayer centre. 24SM-P, DOPC-P, DPPC-P and cholesterol- O3 depict the average location of the phosphate or oxygen atom of the lipid head groups. 24SM-C18, 24SM-C24, DPPC-C16 and DOPC-C18 depict the location of the lipid acyl chains. The locations of only the terminal acyl chain carbons are displayed to demonstrate the depth of acyl chain penetration into the bilayer. **B**) (***a***) Free energy profile for transferring a cholesterol molecule from outer leaflet to the inner leaflet in a C16 SM asymmetric membrane shows cholesterol has a slight preference for the outer leaflet. The Z-axis refers to the proximity of the lipids relative to the bilayer centre (see Methods). (***b***) Normalized density profile for specific atoms of different lipids in the 16:0 SM membrane system. The Z-axis refers to the proximity of the lipids relative to the bilayer centre. 16SM-P, DOPC-P, DPPC-P and cholesterol-O3 depict the average location of the phosphate or oxygen atom of the lipid head groups. 16SM-C18, 16SM-C16, DPPC-C16 and DOPC-C18 depict the location of the lipid acyl chains. The locations of only the terminal acyl chain carbons are displayed to demonstrate the depth of penetration into the bilayer.

**Supplementary Fig. 8.**
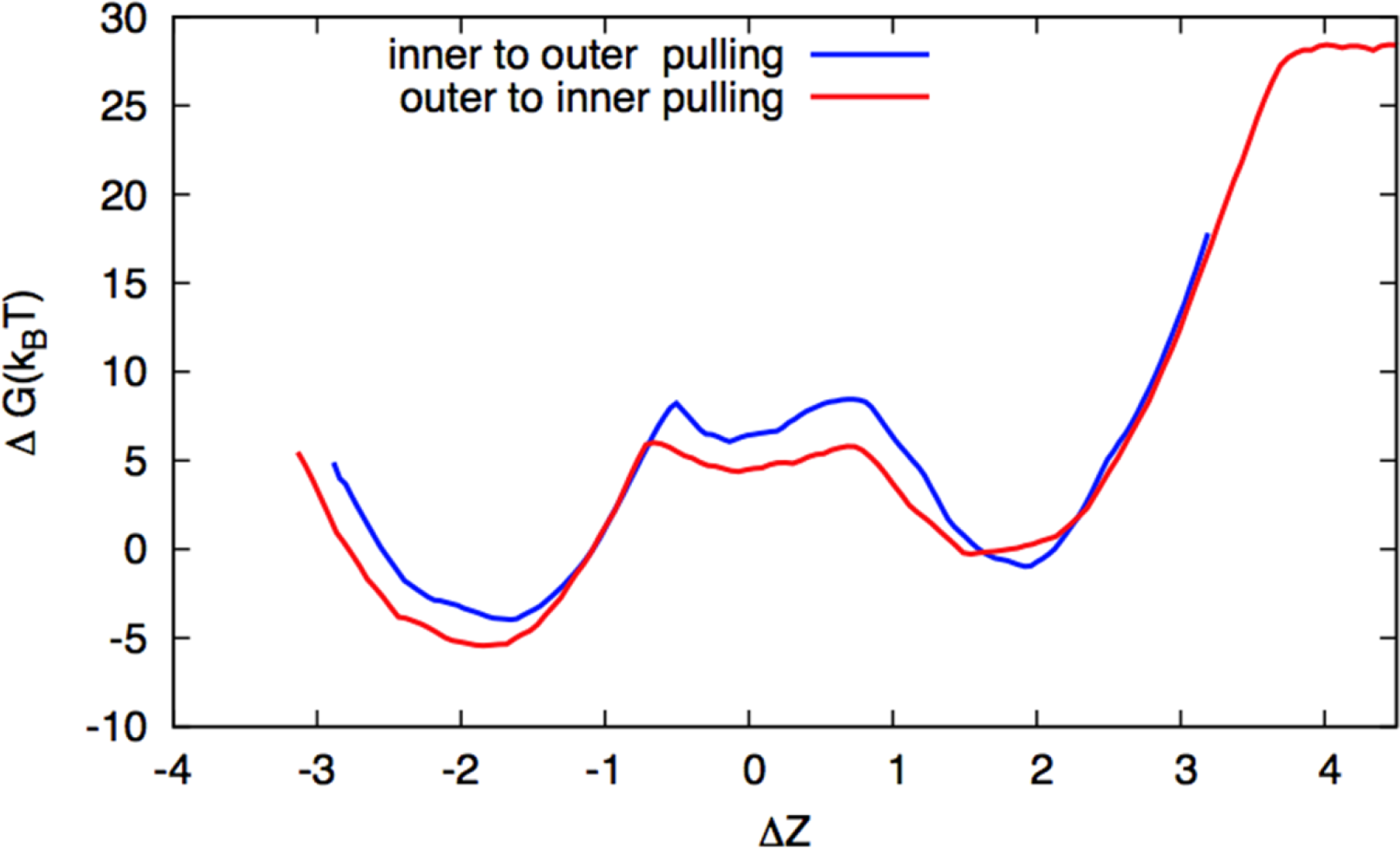
Free energy of cholesterol movement across the bilayer is unaffected by the directionality of the Potential of Mean Force (PMF) calculation. Free energy profiles for the asymmetric C24 SM systems when cholesterol was pulled from the outer to inner (red) and inner to outer (blue) monolayers. The same trend (cholesterol favouring the inner leaflet) is observed, regardless of PMF directionality. The error bars are not shown to clearly distinguish between the two free energy profiles.

**Supplementary Fig. 9.**
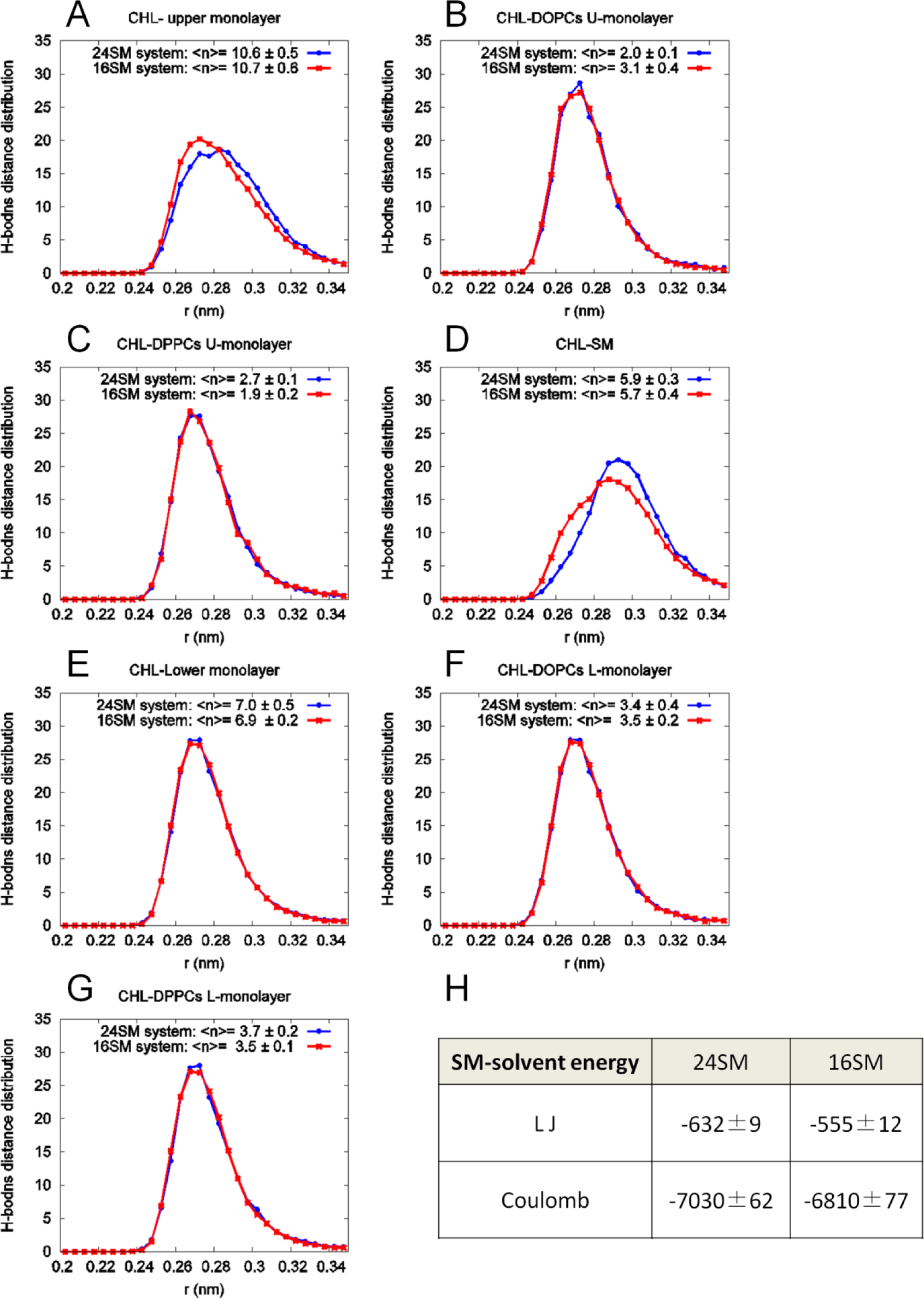
Cholesterol exhibits reduced hydrogen bonding strength with C24 sphingolipids compared to C16 sphingolipids in asymmetric membrane molecular dynamic simulations. Each plot is a distribution of the H-bond distances formed between specific species: **A)** Cholesterol and all upper leaflet lipids. **B)** Cholesterol and upper leaflet DOPC. **C)** Cholesterol and upper leaflet DPPC. **D)** Cholesterol and upper leaflet SM. **E)** Cholesterol and all lower leaflet lipids. **F)** Cholesterol and lower leaflet DOPC. **G)** Cholesterol and lower leaflet DPPC. The <n> value in the upper right corner of each panel refers to the number of H-bonds made by cholesterol in each plot. The data (A and D) shows that the H-bonding with 16 SM occurs at shorter distances (corresponding to a stronger H-bond) than with 24 SM. All other distributions are identical. **H)** Comparison of Lennard-Jones (LJ) and electrostatic interaction energies between C16 or C24 SM and the solvent molecules.

**Supplementary Fig. 10.**
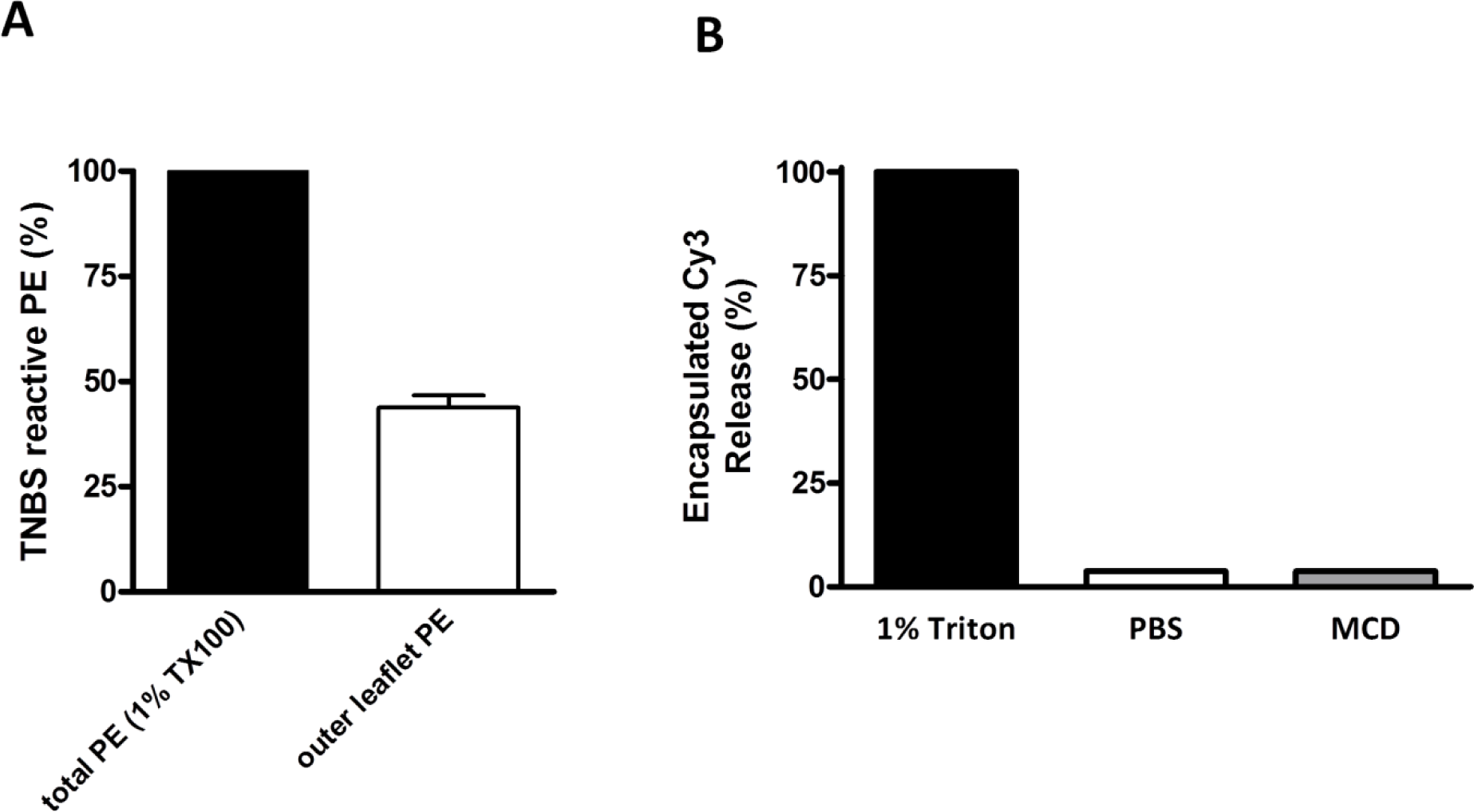
Vesicles are unilamellar and remain intact after cholesterol extraction with cyclodextrin. **A)** Symmetric LUVs containing phosphatidylethanolamine (PE) were incubated with membrane impermeable trinitrobenzylsolphate (TNBS). Only outer leaflet PE will react with TNBS. The amount of reactive PE in intact vesicles was determined by reading absorbance at 410nm compared with TNBS reactive PE from TX-100 solubilized LUVs (100%). TNBS modified 50% PE of LUVs, demonstrating that PE is equally distributed between leaflets and that the LUVs are unilamellar. **B**) LUVs with encapsulated Cy3 were treated with PBS, MCD (5 mM) or dissolved with 1% TX-100. The amount of Cy3 released into the media after 30 minute treatment was quantified by fluorescence spectroscopy. MCD treatment did not cause Cy3 leakage and thus did not affect LUV structural integrity.

**Supplementary Fig. 11.**
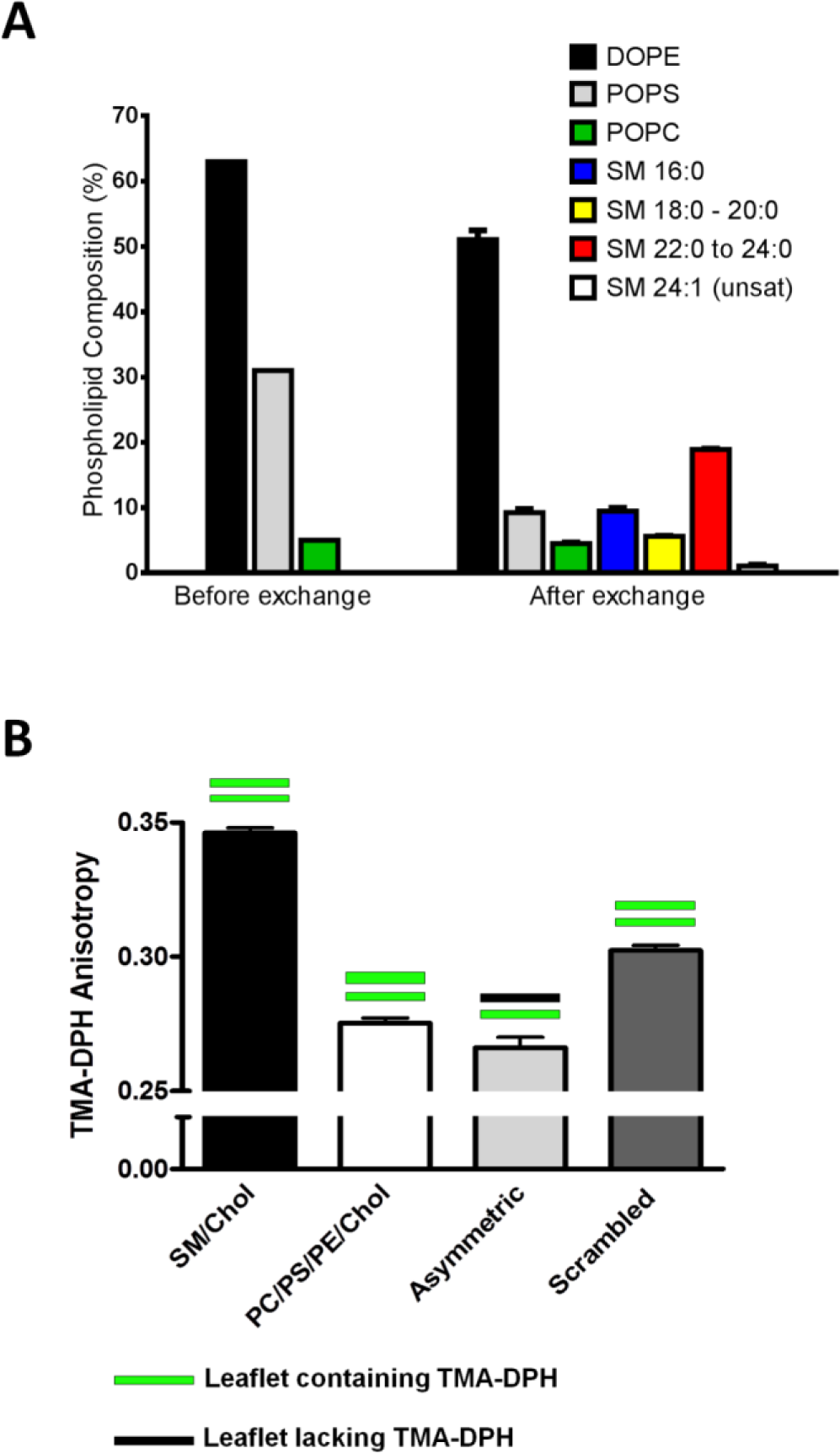
Characterization of transversely asymmetric large unilamellar vesicles. **A)** Representative phospholipid composition in acceptor PE/PS/PC LUVs before and after outer leaflet exchange with milk SM LUVs by mass spectrometry analysis. **B)** LUVs were made with 1% TMA-DPH. SM/cholesterol LUVs (black bar) had higher anisotropy value than PC/PS/PE/cholesterol LUVs (white bar). Asymmetric vesicles were then generated from these PC/PS/PE/cholesterol LUVs by changing with C24 SM/cholesterol LUVs that contain no TMA-DPH. Resulting asymmetric LUVs maintained the acceptor’s anisotropy, as TMA-DPH remained in the inner leaflet after the outer leaflet exchange with SM (asymmetric, light gray bar). However, if the same LUVs were dissolved and remade into symmetric LUVs (scrambled, dark gray bar), anisotropy took an intermediate value between donor and acceptor. TMA-DPH now senses the effect of SM in the LUVs, indicating the successful introduction of SM into the outer leaflet of the acceptor LUVs.

**Supplementary Fig. 12.**
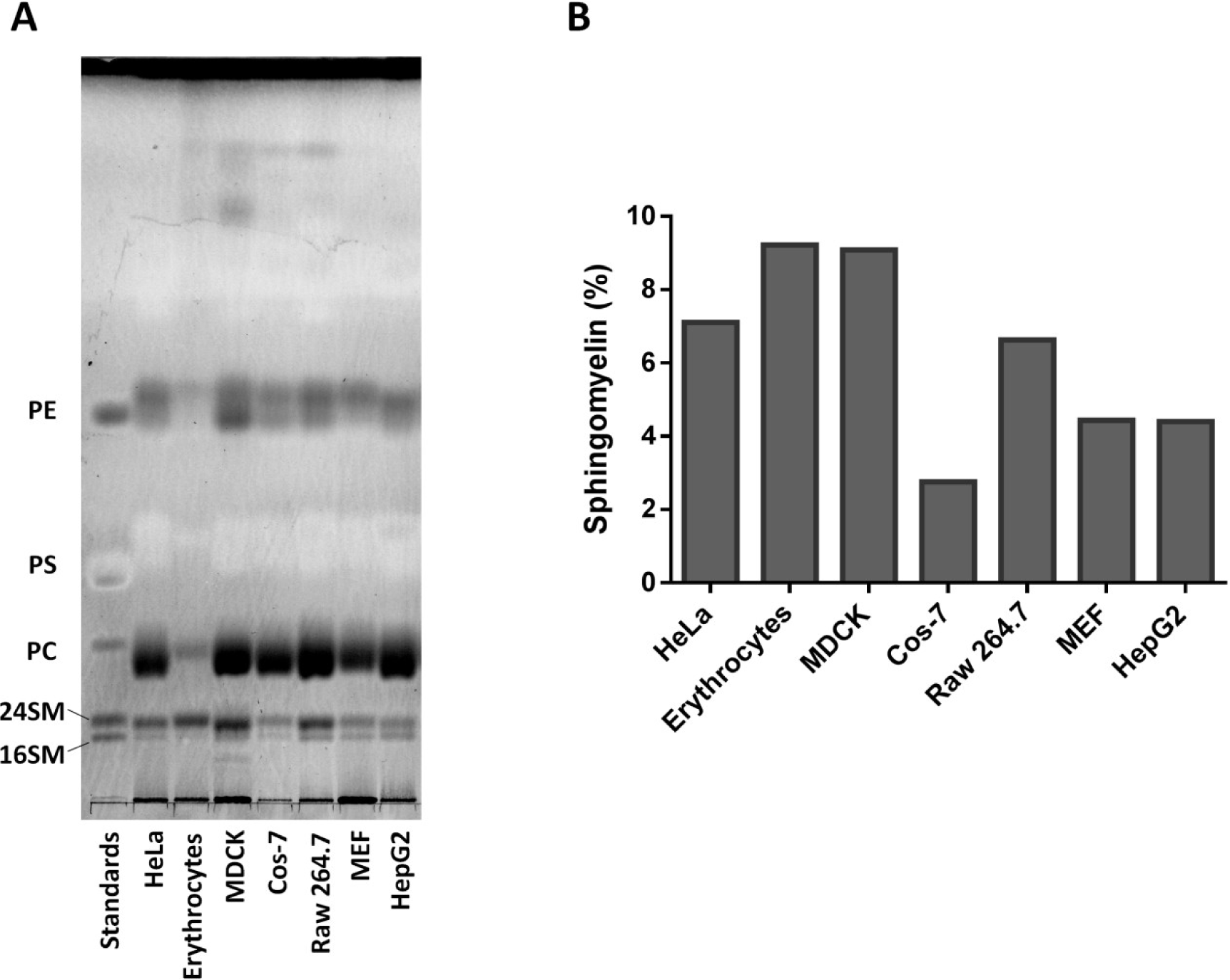
C24 sphingolipids are abundant in many cell types. **A**) Total lipids were extracted from multiple mammalian cell types and resolved by thin layer chromatography (TLC) using a chloroform/acetone/methanol/acetic acid/water (6:8:2:2:1) solvent system. Lipids on the plate were visualized with coomassie blue G stain. **B**) Densitometric quantification of TLC sphingolipid content, relative to total lipid staining.

**Supplementary Fig. 13.**
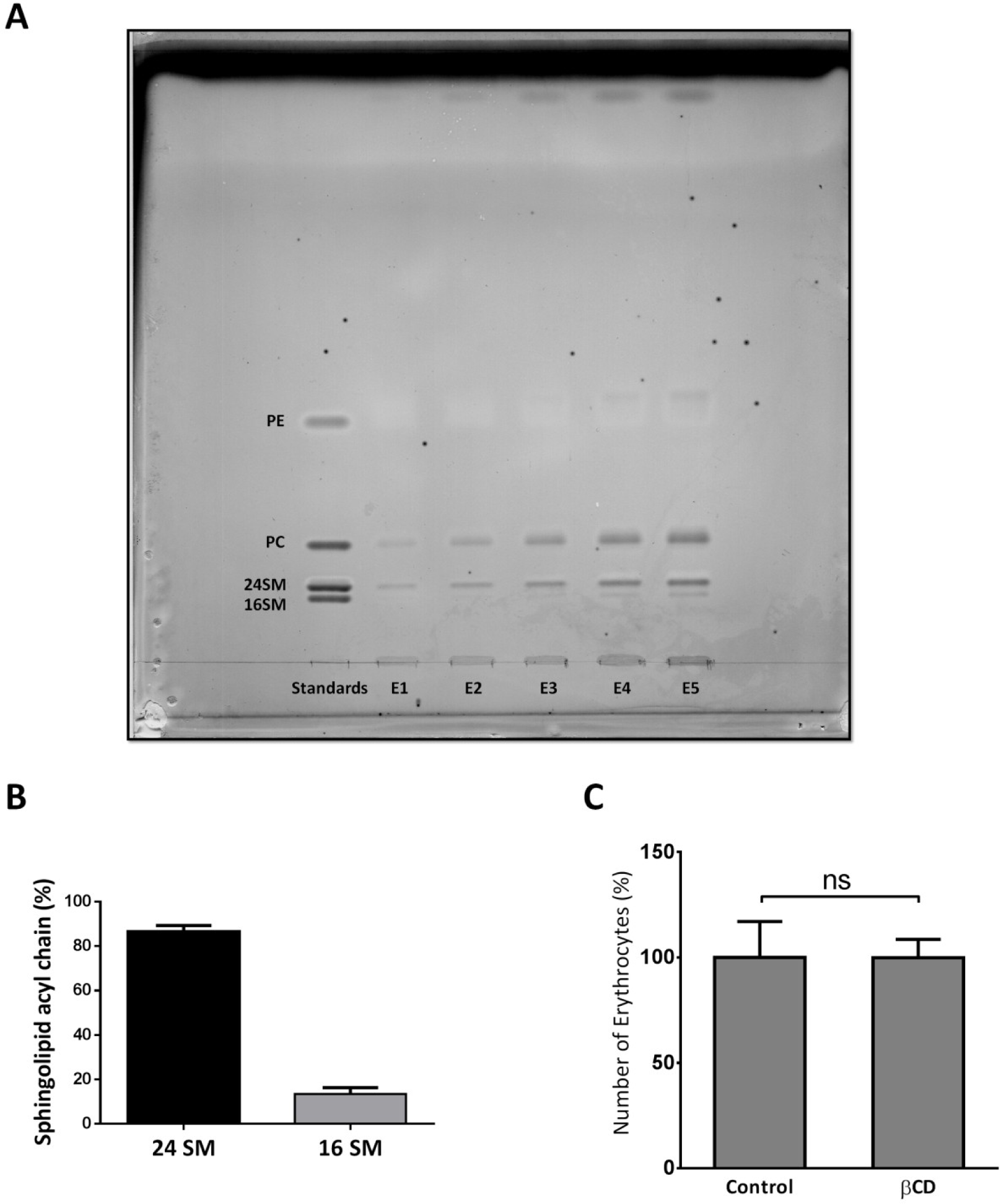
Human erythrocytes contain predominantly C24 acyl chain sphingolipid and erythrocytes remain intact during cholesterol exchange protocol. **A**) Representative thin layer chromatography showing the relative quantities of lipids isolated from human erythrocytes. Lane 1, lipid standards. Lanes 2-6 show increasing quantities (E1-60 μL, E2-90 μL, E3-130 μL, E4-170 μL, E5-200 μL) of lipids isolated from human erythrocytes. **B**) Quantification of the relative abundance of C16 and C24 sphingolipids isolated from human erythrocytes. **C**) Quantification of erythrocyte number after exchange protocol with medium along or plus 1mM βCD. Error bars represent standard error of the mean from 3 independent experiments.

### Materials and Methods

#### Cell culture

HeLa cells were cultured in Dulbecco’s Modified Eagle Medium (DMEM) supplemented with 10% fetal bovine serum and 1 % antibiotics (penicillin and streptomycin) at 37 °C with 5 % CO_2_. For some experiments, the culture medium was changed to serum-free DMEM supplemented with 1 mg/mL bovine serum albumin overnight.

#### GUV preparation

Symmetric GUVs containing 0.05 % rhodamine-DPPE were created by depositing 2.5 µL of 330 µM lipids dissolved in chloroform:methanol (95:5) onto two 3 cm long platinum wires positioned 3 mm apart (41). Organic solvent was evaporated away under a stream of nitrogen gas and then the wires were placed in a vacuum desiccator for at least 1 hour. The platinum wire was then submerged into approximately 1 mL of 30 mM sucrose solution at 70 ºC and connected to a digital PM5193 programmable synthesizer/ function generator producing an A/C sine wave at 10 Hz and 3 V for 90 minutes. For symmetric GUVs, the vesicles also contained 0.1 % NBD-DPPE. For microscopy purposes, the sucrose encapsulated vesicles were diluted into 30 mM glucose solution in order to settle the vesicles onto the bottom of a microscope dish.

#### Asymmetric GUVs

Symmetric GUVs were initially generated and converted into asymmetric GUVs by outer leaflet lipid exchange by adapting a previously published protocol (17). 5 mM cholesterol-containing outer leaflet donor LUV lipids plus 0.1 % NBD-DPPE were generated in 30mM sucrose and incubated while vortexing for two hours at 55 ºC in 60 mM HPα-CD. DPPC/DOPC/Cholesterol (35:35:30) GUVs containing 0.05 % rhodamine-DPPE were then incubated in 20 mM of the HPα-CD/ donor lipid complex at 70 ºC for 30 minutes. To remove excess donor lipid and HPα-CD, the samples were washed in 30 mM glucose by passively filtering away the donor LUVs through a membrane with 8 µm pores and retaining the asymmetric GUVs. The resulting asymmetric GUVs were then transferred to a microscope dish.

#### LUV preparation

All lipids were stored as 50 mg/mL stocks in chloroform/methanol (95:5). 1 mM LUVs are generated by combining the desired lipid components followed by evaporating the organic solvent under nitrogen gas and subsequent vacuum desiccation for at least 60 minutes. In the case of symmetric ^3^H-cholesterol labeled LUVs, trace amount of ^3^H-cholesterol (~100 pmol/3 µmol phospholipid) was incorporated into the sample prior to desiccation. Dried lipids were then resuspended in Medium 1 (20 mM HEPES, 150 mM NaCl, 5 mM KCl, 1 mM CaCl_2_, 1 mM MgCl_2_), subjected to 5 freeze/heat cycles and extruded through a 100 nm membrane to form 100 nm LUVs. For LUV isolation from the aqueous media, 1 mol% biotinylated-PE was added to lipid mixture above and resultant LUVs were incubated with streptavidin-coated agarose beads at 4 °C overnight. Unbound LUVs were removed by washing the beads 3 times with Medium 1. For intermembrane exchange, two LUV populations (donor and acceptor) were similarly made with 30 % cholesterol. The donor population received trace ^3^H-cholesterol, was biotinylated and bound to streptavidin beads to facilitate separation from unlabelled acceptor LUVs. Before conducting MCD experiments, LUV properties were analyzed to determine the size, integrity and unilamellarity (a single bilayer).

#### Asymmetric LUVs

Asymmetric LUVs with POPC, eSM, bSM or mSM introduced into the outer leaflet and POPC/POPS/POPE (1:1:1) in the inner leaflet were generated by adapting a previously published protocol (19). Experiments were also conducted using DOPE, POPS, POPC (2:1:0.15) as the inner leaflet. Briefly, 500 µL of 10 mM donor LUVs (POPC, eSM, bSM or mSM) in Medium 1 were mixed with 100 µL 360 mM hydroxypropyl-α-cyclodextrin (HPα-CD) and vortexed for 2 hours at 55 °C. The donor-HPα-CD solution was then mixed with 600 µL of 2 mM acceptor LUVs containing 25 % sucrose for 30 minutes at 55 °C to initiate outer leaflet lipid exchange. After the mixture had cooled to room temperature, the 1 mL solution was overlaid onto 4 mL 10 % sucrose in Medium 1 and centrifuged at 10 °C in an NVT-100 rotor for 40 minutes at 190,000 x g. After centrifugation, the excess donor-HPα-CD complex remains in suspension, while the sucrose-containing acceptor vesicles are collected as a pellet. Centrifugation of the resulting asymmetric vesicles was repeated a second time followed by a final resuspension in 1 mL Medium 1. When examining the effect of transbilayer asymmetry on cholesterol distribution, the 10 mM LUV donor contained 1% biotinyl-PE allowing for the final asymmetric vesicles to be bound to streptavidin agarose beads. ^3^H-cholesterol was introduced into the asymmetric vesicles by incubation with 1 mM ^3^H-cholesterol donor LUVs in the presence of 1 mM βCD at 37 °C followed by repeated washing to remove unincorporated ^3^H-cholesterol. The cholesterol transbilayer distribution was determined by MCD mediated extraction, as described below.

#### Microscopy

All fluorescence images of GUVs were generated on a Nikon TE2000-E inverted fluorescent microscope with a 60x objective. Data was captured by 100-200 millisecond exposure with a Photometrics Cascade 512B CCD camera and Metamorph software. GUVs were found in the sample by searching the sample under low light conditions or by Differential Interference Contrast microscopy to avoid photo-oxidation effects. Post-acquisition editing was performed using ImageJ v1.43 software.

#### Molecular dynamics simulations

All-atom MD simulations and PMF calculations was performed using GROMACS version 5.1 (42, 43)and the CHARMM36 force field (FF)(44, 45). The TIP3P solvent model was used(46). Electrostatic interactions were treated with particle mesh Ewald (PME) with a shortrange cutoff 1.2 nm, and van der Waals interactions were switched off between 1.0 to 1.2 nm. For all systems temperature was kept constant at 37 **º**C using Nose-Hoover temperature coupling(47, 48). Bonds containing hydrogen atoms were constrained using the LINCS (49) algorithm. Parrinello-Rahman barostat pressure coupling (50)(tau_p=1 ps) was applied on all systems after equilibrating the systems with Berendsen pressure coupling(51). The leap-frog integrator was used with a timestep of 2 fs. The systems were built using the CHARMM-GUI membrane builder. After a short equilibration for 20ns, each simulation was continued for 500 ns for data sampling.

#### Potential of mean force calculation

PMF calculation was performed to obtain the free energy profile of cholesterol flip-flop across the bilayer normal. The model bilayer for the PMF calculation contained ~ 128 lipids. The lipid concentration was setup carefully, so that the total area of the monolayers matched as closely as possible. System composition for each system is: upper monolayer: SM (18), cholesterol (19), DOPC (15) DPPC (15), lower monolayer: cholesterol (19), DOPC (22), DPPC (23), with 6878 water molecules.

For each system, 40 windows were generated by pulling a cholesterol (target cholesterol) molecule across the bilayer. The distance between the phosphorus atom of a specific SM molecule (reference SM) and Oxygen atom of the target cholesterol was chosen as the reaction coordinate. The maximum distance between reaction coordinates in adjacent windows is around 0.2 nm. The phosphorus atom of the reference SM was restrained in the *z* direction (bilayer normal) by a spring potential with a spring constant of 1000 kJ/mol. The spring constant of the biasing potential is 1000 kJ/mol. Each window was simulated for 100ns, which was sufficient for the target cholesterol to sample the entire XY plane in each simulation window. Pulling a cholesterol molecule in the opposite direction (inner to outer leaflet) did not change the PMF profiles. The windows were generated using the pull option in GROMACS, by restraining the P atoms of the reference SM in the *z* direction. The pulling force (along *z*) was applied on the O atom of the target cholesterol. In addition, to force cholesterol to flip orientation during the pulling (to save computational time) the last carbon atom of the target cholesterol was restrained (in the z direction) by a spring potential (spring constant 100 kJ/mol). This generates a torque when the cholesterol crosses the middle of the bilayer.

#### Temperature controlled cholesterol extraction by MCD

Cholesterol flip-flop in symmetric and asymmetric LUVs was controlled by reducing the temperature to 0 °C. We developed a rather rigid protocol, which we found was absolutely necessary, in order to successfully and reproducibly prevent flip-flop. The temperature of the cholesterol extraction procedure was stringently regulated by performing the experiments in a cold room and in an ice water bath, or 0 °C water bath containing 50 % ethylene glycol. Furthermore, to prevent incidental hand warming of the samples, all manipulation of the tubes was required to be carried out with utensils similarly maintained at 0 °C. Samples, utensils and MCD media were pre-incubated at 0 °C at least one hour prior to the cholesterol extraction. Without sufficient 0 °C cooling, even utensils used to rapidly transfer tubes, cholesterol flipping would occur (>50% extraction). To initiate cholesterol extraction, 5 mM MCD was added to the medium to selectively remove cholesterol from the outer leaflet of the LUVs 0 °C until cholesterol extraction reached a plateau. Similar experiments were also conducted in a 37 °C water bath. After incubation, the LUVs were separated from MCD containing medium by a brief centrifugation in 0 °C centrifuge within a cold room. The amounts of ^3^H-cholesterol in the medium and in the LUVs are quantified by scintillation counting. Total ^3^H-cholesterol was determined by lysing the same amount of ^3^H-cholesterol labeled LUVs with 2% Triton X-100.

#### LUV intermembrane exchange

The assay was performed by mixing bead-bound ^3^H-cholesterol donor LUVs with excess unlabelled acceptor LUVs (100-fold) at 37 °C, 0 °C or -5 °C in the presence of 1 mM βCD as shuttle. Outer leaflet cholesterol is exchanged between populations, until equilibrium, at 0 °C or -5 °C. Donor and acceptor LUVs are isolated by 60 second centrifugation at 1000 x g and the amount of ^3^H-cholesterol in the acceptor LUVs is quantified.

#### Lipid Mass Spectrometry

Confirmation of outer leaflet lipid exchange in LUVs was performed by electrospray ionization tandem *mass spectrometry* (ESI-MS/MS). Asymmetric LUVs were generated as described above. The lipids were then extracted by the Bligh and Dyer method (52) in the presence of internal standards including 1,2-dieicosanoyl-sn-glycero-3-phosphocholine (PC), N-heptadecanoyl-sphingomyelin (SM), 1,2-ditetradecanoyl-sn-glycero-3-phosphoethanolamine (PE), and 1,2-ditetradecanoyl-sn-glycero-3-phosphoserine (PS). Lipid extracts from LUVs were then subjected to shotgun lipidomics in the negative ion mode using neutral loss scanning (NLS) for 50 amu (for PC), NLS for 87 amu (for PS), and product ion scanning for m/z 196 (for PE). Individual lipid molecular species were quantified by comparing the ion intensity of individual molecular species to that of the lipid class internal standard following corrections for type I and type II ^13^C isotope effects.

#### Asymmetric vesicle TMA-DPH anisotropy

Fluorescence anisotropy measurements were performed using a Photon Technology International fluorometer and FeliX software. We generated symmetric SM/cholesterol and PC/PS/PE/cholesterol vesicles as described previously with the addition of 1 mol% TMA-DPH. To generated asymmetric vesicles, SM/cholesterol was introduced into the outer leaflet of PC/PS/PE/cholesterol vesicles. During the asymmetry procedure no TMA-DPH was present in the SM/cholesterol donor vesicles. Outer leaflet TMA-DPH was then removed from PC/PS/PE/cholesterol vesicles during phospholipid exchange resulting in only inner leaflet labeling of asymmetric vesicles. Anisotropy was compared between symmetric, asymmetric and scrambled vesicles by excitation and emission at 365 nm and 425 nm, respectively.

#### Erythrocyte intermembrane exchange

All erythrocyte work was approved by the Ottawa Health Sciences Network Research Ethics Board (#20140233-01H). Human erythrocytes were isolated from whole blood by centrifugation through Ficoll-Hypaque gradient solution. The erythrocytes were then washed 3 times with PBS containing 2 mM EDTA. 100 million washed erythrocytes were incubated with 20 µCi ^3^H-cholesterol in 10 mL DMEM for at least 4 hours, subsequently biotinylated according to manufacturer’s instructions (Fisher Scientific) and 100,000 cells were adhered to a streptavidin coated microplate. 100-fold excess unlabelled erythrocytes were added to the microplate wells in suspension with 1 mM βCD at 37 °C or 0 °C, until equilibrium. 75 µL of supernatant containing erythrocyte acceptor cells was removed at each time point and the amount of cholesterol transferred to the acceptors was determined by scintillation counting.

#### Fluorescence resonance energy transfer efficiency simulation

A Monte Carlo approach was used for simulations of density-dependent FRET using a program written in Fortran 95 and compiled using gfortran/gcc running in the command line interface of a computer running Mac OSX 10.8. The number of YFP and CFP molecules per 1000 nm X 1000 nm square were input. YFP and CFP molecules were assigned floating-point coordinates at random within a square patch of simulated membrane using the Mersanne Twister pseudorandom number generator and units of nm. Molecules were not allowed to fall within 2.4 nm of other molecules, as this is the diameter of the GFP cylinder. Once molecules were placed, FRET efficiency (E) was calculated for each CFP to every YFP within 50 nm of the CFP using the equation E = 1 / (1 + (r/R_0_)^6^ where r is the distance between the centers of the two molecules and R_0_ is 4.89 nm (53). At greater than 50 nm, E < 1 × 10^−6^, and FRET was not calculated for reasons of computational efficiency. Cumulative FRET efficiency from a CFP was calculated from the cumulative probability of FRET from this CFP molecule to all available YFPs.

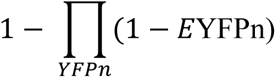

Total FRET efficiency was calculated as the mean efficiency of FRET from all CFPs included in the simulation.

#### Live cell fluorescence resonance energy transfer

Fluorescence resonance energy transfer (FRET) was performed on HeLa cells co-transfected with mCFP- and mYFP-GPI-APs. Images were acquired with a Nikon TE2000-E inverted fluorescent microscope using a 60x objective. Data was captured on a Cascade 512B CCD camera (Photometrics) and MetaMorph software (Universal Imaging). To quantify the crosstalk between CFP and YFP channels, cells were transfected with either mCFP or mYFP plasmids and imaged identically as in FRET experiments. This generates crosstalk factors from CFP or YFP to the sensitized YFP channel, G_CFP_ and G_YFP_, respectively. To measure the FRET, co-transfected cells were imaged with a 3-cube system: CFP_ex_/CFP_em_ (I_CFP_), CFP_ex_/YFP_em_ (I_S_) and YFP_ex_/YFP_em_(I_YFP_). The true FRET signal, I_FRET_, was calculated as following (54):

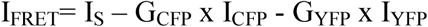

FRET efficiency, E (%), was derived as below:

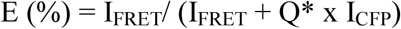

*Q is the ratio of sensitized emission, I_FRET_, to the corresponding amount of donor (CFP) recovery in CFP_ex_/CFP_em_ channel after YFP photobleaching measured in the same cell. Q was determined experimentally using co-transfected HeLa cells. Images were acquired using the same 3-cube system and imaging parameters as the experimental images, and FRET efficiency was determined afterward for the same cells using the acceptor photobleach method (54).

The final E (%) data was presented after excluding outliers from the normal distribution (mean ± 95% confidence interval).

The protein density of mCFP- and mYFP-GPI-AP in the HeLa cell plasma membrane was determined by imaging purified mCFP and mYFP protein in solution under identical imaging conditions as the FRET experiments. Serial dilutions of the fluorescent proteins in a chamber of 50 μm thickness were imaged to produce a standard curve for fluorescent intensity as a function of protein concentration. The imaging volume was determined using the known pixel dimensions (xy) of the Cascade 512B CCD camera and by placing the fluorescent protein solutions in 50 μm thickness chambers to standardize the focal depth (z). The standard curves for the soluble fluorescent protein densities were then used to interpolate the mCFP- and mYFP-GPI-AP protein densities from the FRET experiments.

For sphingomyelin manipulation, the mCFP- and mYFP-GPI-APs expressing HeLa cells were treated with or without 50 μM myriocin and fumonisin B1 for 3 days to deplete the sphingolipids. The night before microscopy experiments, the media was changed to serum free media to prevent incorporation of exogenous lipids from the fatal bovine serum into the cells. In the morning, a subset of the cells was incubated with either 16:0 SM or 24:0 SM/γ-CD complexes (20µM SM and 1mM γ-cyclodextrin) for 1 hour at 37 °C. Microscopy experiments were performed at approximately 12 °C. Each treatment was also accompanied by an additional dish treated with 0.2 % saponin for 30 minutes on ice before microscopy experiments. This was to compare the FRET change before and after depletion of cholesterol for each treatment.

#### Sphingolipid analysis

To confirm thorough depletion of sphingolipids by the myriocin and fumonisin b1 treatment, as well as, to test the efficiency of sphingomyelin supplementation, thin layer chromatography (TLC) was performed on lipid extracts from HeLa cells. After each treatment, HeLa cells were trypsinized and washed in PBS, followed by lipid extraction (52). Lipid extracts were loaded onto the TLC plate alongside purified lipid standards to compare sphingolipid levels in all treatments. The lipids were resolved on the plate by running in a chloroform/acetone/methanol/acetic acid/water (6:8:2:2:1) solvent system. The lipids were visualized by incubating the plate in a solution of 0.03 % coomassie blue G, 30% methanol and 100 mM NaCl followed by destaining in 30% methanol and 100 mM NaCl. Sphingolipid content was quantified by densitometry analysis using Image J software.

#### Sensitivity to sphingomyelinase

To validate that the re-supplied sphingolipids were successfully incorporated into the plasma membrane of the HeLa cells, we quantified sphingomyelinase-induced endocytosis of FITC-dextran (25). HeLa cells were treated either with DMSO (control) or Fumonisin B1 (50µM) and Myriocin (50µM) in DMEM containing 1mg/ml BSA for 72 hours. Post drug treatment, C16 or C24 SM were added back to the cells as γ-cyclodextrin complexes. SM (20µM) and γ-cyclodextrin (1mM) complexes were added for 1 hour at 37 °C. To visualize endocytosis, FITC-dextran (5mg/ml) and sphingomyelinase (50mU/ml) were added to the cells for 10 minutes at 37 °C. The cells were then washed in PBS, fixed in 4% paraformaldehyde and mounted with slow-fade DAPI. Fluorescence microscopy was performed using Zeiss LSM 510 confocal microscope. Intensity of FITC-dextran was measured using Image J and normalized to the DMSO control. Error bars represent standard error of the mean from three independent experiments.

#### Statistical analysis

Statistical differences in cholesterol transbilayer distribution between different asymmetric LUVs were determined by one-way ANOVA. Post hoc comparisons were conducted relative to the mSM asymmetric sample with * indicating P<0.05, ** indicating P<0.01 and *** indicating P<0.001. For comparisons of erythrocyte number with and without cyclodextrin treatment, statistical differences were examined using an unpaired Student’s *t*-test. Non-linear regressions were performed in Graphpad Prism 5.0 and fit using the equation: Y=Top*(1-exp(-K*X)). Error bars throughout represent standard error of the mean from 3 independent experiments.

## Footnotes

* DPPC: dipalmitoyl-phosphatidylcholine; DOPC: dioleoyl-phosphatidylcholine; DPPE: dipalmitoyl-phospatidylethanolamine

